# Memory genes respond strongly to genetic perturbations and have CpG-depleted promoters

**DOI:** 10.64898/2026.11.28.769752

**Authors:** Ida Büschel, Vivienne Nahrstedt, Benjamin Kuznets-Speck, Carsten Marr, Yogesh Goyal

## Abstract

Genetically identical cells in the same environment differ in gene expression. For some genes, differences persist across cell divisions. Available methods identify such memory genes using different statistics and have never been compared on the same cells. What distinguishes memory genes, and how they act within a regulatory network, is unknown. Here we show that four methods converge on a common set of memory genes, which are characterized by slow turnover, with longer mRNA half-lives and CpG-depleted promoters, but not by more frequent or longer bursts. Across 21 CRISPR screens covering 78,210 perturbations and 22 million cells, these genes responded more strongly to perturbation than expression-matched controls, while perturbing them produced no larger response than perturbing any other gene. A theoretical model, validated empirically across datasets, traces their stability and responsiveness to a single relaxation rate. Expression memory therefore has a hitherto unappreciated function: predicting how strongly a gene responds to perturbation.

## Introduction

Genetically homogenous cells in the same environment can differ markedly in gene expression, and some of these differences can persist across cell divisions. Such heritability has been observed for single proteins and transcripts, and for high-dimensional cell states, lasting several generations^1–4^. This “memory” can have functional consequences. Rare, transiently persistent states in drug-naive cancer cells seed therapy resistance^5–9^, determine which somatic cells reprogram to pluripotency^10,11^, and set susceptibility to viral infection^12,13^. How memory arises at the transcriptional level is only partly understood. Network models with transcriptional bursting reproduce these states^14^, and chromatin-level mechanisms can write transient responses into lasting memory^15^. However, which genes show heritable expression (henceforth “memory genes”) has only recently become measurable at genome scale.

Memory genes are defined as genes whose expression is more similar among clonally related cells than among unrelated cells of the same population (**Fig. 1a**). Memory genes were first identified with MemorySeq^16^, by expanding single cells into clones and profiling each in bulk: genes whose state is inherited vary between clones, whereas fast-switching genes do not. Since then, we and others have combined lineage barcoding with single-cell RNA sequencing to compare gene expression between clonally related cells and unrelated cells from the same population^7,15–19^. Several methods now identify memory genes from such data: by the fraction of expression variance explained by clone identity^20^, by the variability of clone-level expression^21^, or by learned representations of lineage structure^18^. Others infer memory without experimental lineage barcodes, from the covariance structure of a single expression snapshot^22^ or from lineages predicted from expression^21^. These methods use different statistical approaches to identify memory genes and have not been systematically compared on the same cells. As such, whether they recover a common set of memory genes is unknown. Moreover, memory genes have not yet been studied beyond their association with traits such as drug resistance. We therefore set out to test how they function within regulatory circuits and whether they possess defining genomic, transcriptomic, or epigenetic features.

**Fig. 1:**
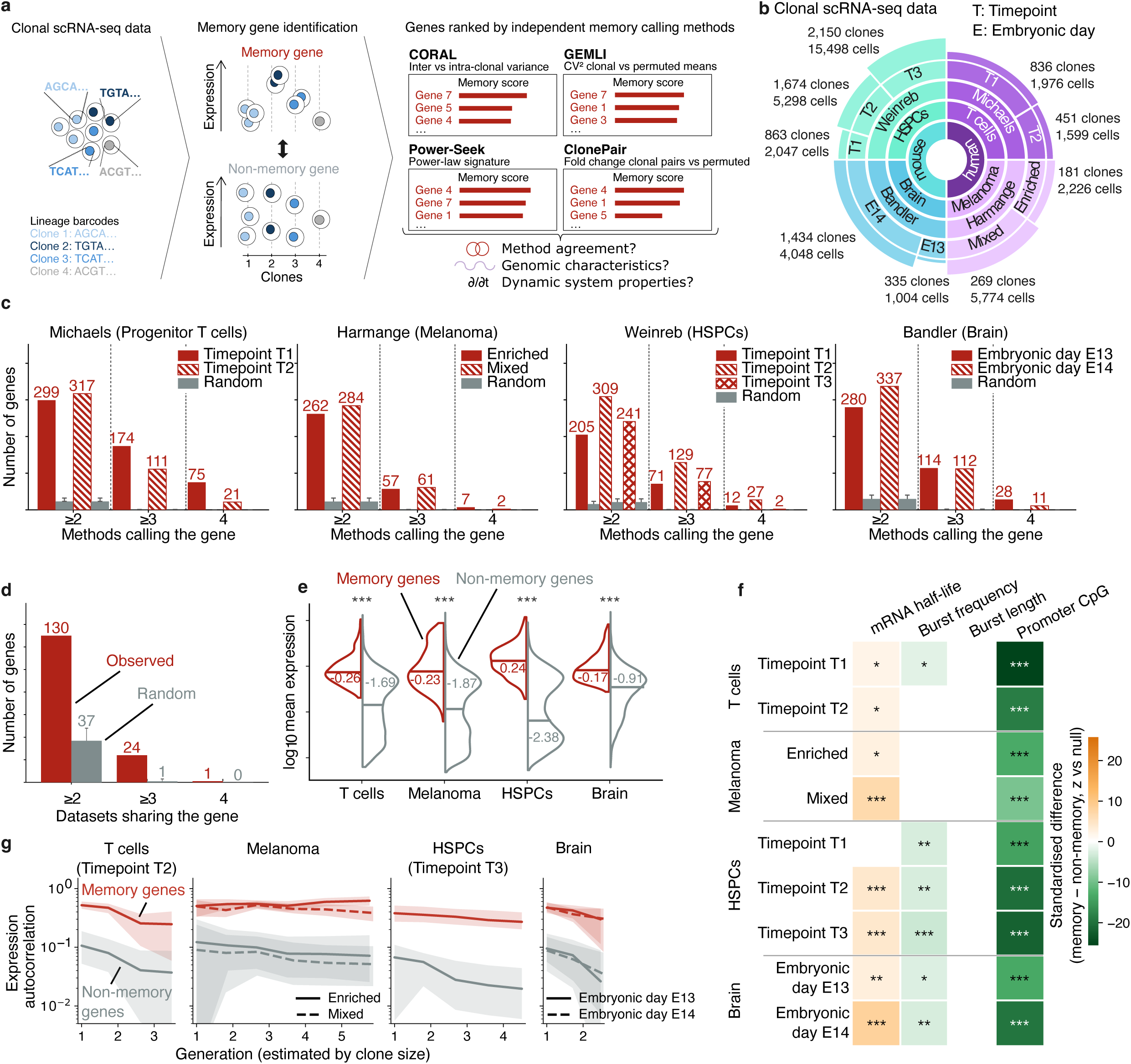
Independent methods converge on consensus memory genes with high expression, long-lived mRNAs, CpG-depleted promoters, and persistent expression autocorrelation. **a** Four methods rank every measured gene by a different memory statistic. Left, clonally resolved scRNA-seq: cells sharing a lineage barcode form a clone. Centre, the distinction every method makes: memory genes are homogeneously expressed within clones, whereas non-memory genes vary as much within clones as across the population. Right, the four memory-calling methods and the statistic each uses: CORAL^15^, inter-versus intra-clonal variance; GEMLI^20^, squared coefficient of variation of clonal versus permuted means; Power-Seek^21^, power-law eigenspectrum; ClonePair, fold change in clonal versus permuted cell pairs. Each method returns all measured genes ranked by its own memory score. **b** Memory genes are identified in four clonal scRNA-seq datasets with 6,863 distinct clones and 39,470 cells in total. Rings from the centre: species (human, mouse), tissue (T cells, hematopoietic stem and progenitor cells, brain, melanoma), dataset (first author), experimental group: differentiation timepoints (Michaels^22^, timepoints 1, 2; Weinreb^23^, timepoints 1 to 3), sorted states (Harmange^18^, enriched and mixed), or embryonic day of barcode injection (Bandler^24^, days 13, 14). Each dataset occupies a quarter of the circle. Within a dataset, wedge width is proportional to the number of clones with ≥2 measured cells in each group, and labels give clone and cell counts. Clones with only a single measured cell are excluded throughout. **c** Memory genes called by two or more methods, per dataset and group. Coloured bars, number of genes ranked among the top 300 by at least two, at least three, or all four methods, one bar per experimental group; numbers above bars, observed gene counts. Grey bars, the permutation null, in which each method independently draws the same number of genes without replacement from all measured genes (2,000 permutations; bar height, permutation mean; whisker, 97.5th percentile). Genes called by at least two methods, merged across a dataset’s experimental groups, define that dataset’s consensus memory genes (**Methods**). **d** Consensus memory genes identified in two or more datasets. Red bars, the number of consensus memory genes shared by at least two, at least three, or all four datasets; grey bars, an expression-matched null in which each dataset’s gene set is randomly redrawn within its own expression-rank bins, preserving set size and expression profile (1,000 permutations; bar height, mean; whisker, 97.5th percentile). Mouse genes (Weinreb, Bandler) were mapped to human symbols by orthology, genes without an orthologue were dropped, and all sets were restricted to the 13,162 orthologs measured in all four datasets. **e** Distribution of log_10_ mean expression for consensus memory genes (red) and all other measured genes (grey), per dataset. Split violins; horizontal line and number, median of each group; asterisks, two-sided Mann-Whitney U test (***, p < 0.001). **f** Standardised difference between consensus memory genes and non-memory genes four molecular properties, controlled for expression. Rows, dataset and group; columns, mRNA half-life^35^, burst frequency and length^37^, and promoter CpG observed/expected ratio^36^; genes without a value for a given feature are excluded from that column. The test statistic is the difference in mean feature value between the two sets. Memory labels were permuted among genes within expression bins, preserving the number of memory genes per bin, 10,000 times. Colour, z = (observed difference − null mean) / null SD: orange, higher in memory genes; green, lower; white, not significantly different to the null. For promoter CpG observed/expected, green indicates stronger CpG depletion in memory genes. Asterisks, empirical two-sided permutation p-values (*, p < 0.05; **, p < 0.01; ***, p < 0.001). **g** Expression autocorrelation across generations, measured as the intraclass correlation coefficient (ICC) among cells of the same clone, computed per gene for clones binned by size (2, 3-4, 5-8, 9-16, 17-32 and ≥33 cells; one-cell clones excluded, bins with fewer than 15 clones omitted). Since a clone of n cells has undergone ≈log_2_(n) divisions, the mean log_2_ clone size of each bin estimates the number of generations separating its cells (x-axis). Lines, median ICC across consensus memory genes (red) and non-memory genes (grey); shading, interquartile range; line style, experimental group within a dataset. For Michaels and Weinreb, only the group corresponding to the last measured timepoint is shown, as the other groups consist predominantly of clones with two measured cells.

Here, we compared four memory-calling methods, three published^20–22^ and one developed in this study, on four lineage-barcoded single-cell datasets^17,23–25^ spanning human and mouse, development and cancer, comprising 39,470 cells in 6,863 clones. Although the methods apply different statistical approaches, they converged on a shared set of memory genes, with a median of 13 times more than expected by chance. A subset of these memory genes recurred across cell types and species. Memory genes had longer mRNA half-lives and more CpG-depleted promoters than expression-matched non-memory genes, but did not differ in transcriptional burst frequency or burst length. Their expression stayed correlated within clones over several divisions.

We reasoned that pooled CRISPR screens such as Perturb-seq, which measure the transcriptome-wide effects of thousands of genetic perturbations at the single-cell level^26–29^, could reveal how memory genes function within regulatory circuits: whether they control the expression of other genes, or respond strongly when other genes change. We paired each lineage dataset^17,23–25^ with Perturb-seq screens^28–33^ in the matching cell type, covering six screens and 9,551 perturbations with close to 8.7 million cells. Perturbing memory genes did not produce larger transcriptome-wide responses than perturbing non-memory genes. By contrast, across all methods and pairings, memory genes responded more strongly to perturbations than expression-matched non-memory genes. The effect size became larger as memory genes were restricted to those corroborated by multiple methods. This stronger response is predicted by theory: we formulated a mathematical perturbation-response model, verified its underlying assumptions empirically on the perturbation data, and successfully tested its predictions. Our work reveals a fundamental link between a gene’s expression memory and how strongly it responds to perturbation.

## Results

To find out whether different methods identify the same memory genes, we assembled four published lineage-barcoded scRNA-seq datasets spanning human and mouse, development and cancer: human T-cell progenitors^23^, mouse hematopoietic stem and progenitor cells (HSPCs)^24^, mouse brain^25^, and human melanoma^17^, comprising 6,863 clones and 39,470 single cell transcripts (**Fig. 1b**, **Methods**). Clones with a single measured cell allow no within-clone comparison and were excluded. Within each dataset, we analyzed experimental groups (differentiation timepoints, sorted cell states, the embryonic day of barcode injection) separately since they differ in clone size and number (**Fig. 1b**). To each group we applied four methods which score genes on different statistics (**Fig. 1a**): CORAL partitions a gene’s expression variance into inter-and intra-clonal components^20^; GEMLI compares the squared coefficient of variation of clone means with that of “random clones” of the same sizes by permuting clone labels^21^; Power-Seek ranks genes by their power-law signature in the covariance structure of the gene expression matrix, without using clone labels^22^; and ClonePair, developed by us, scores each gene by the fold change in clonal cell pairs over the fold change in permuted pairs (**Methods**). ClonePair is restricted to clones of ≤ 8 cells, keeping the comparison to closely related cells, and permuted pairs are redrawn 100 times. Genes are called when their concordance frequency (i.e. the fraction of draws in which clonal pairs are significantly more similar than permuted pairs) is ≥ 0.8. The method is insensitive to this threshold over a broad range, and the underlying concordance frequency is strongly bimodal, so genes separate cleanly into called and uncalled (**Supplementary Fig. 1a, b**). We compared the four methods on the top 300 genes per method and group, because the methods rank different numbers of genes (**Supplementary Fig. 1c**). Where a method ranked fewer, we used all the genes it returned (e.g., ClonePair at Weinreb^24^ timepoint 1 identified 143 genes), and the permutation null was drawn with the same per-method set sizes. Results were robust to the number of top genes considered (**Supplementary Fig. 1d**).

The methods independently converged on a shared set of memory genes. In every dataset and group, between 205 and 337 genes were ranked in the top 300 by at least two methods (**Fig. 1c**). This is 9 to 15 times (median 13) the overlap expected under a permutation null, in which each method independently draws the same number of genes at random from all measured genes. Genes called by at least three methods (57 to 174 genes) were likewise highly significant, and genes called by all four methods (2 to 75 genes), a level at which the permutation null is approximately zero, even more so.

We next asked whether the same genes carry memory in different tissues and species. Mouse genes were mapped to human symbols by one-to-one orthology^34^, and all gene sets were restricted to the 13,162 orthologs measured in all four datasets (**Supplementary Fig. 1e**). Consensus memory genes, defined as genes ranked in the top 300 by at least two methods, recurred across datasets beyond an expression-matched null: 130 genes were shared by at least two datasets against 37 expected, and 24 by at least three datasets against <1 expected (**Fig. 1d**). This recurrence was not an artifact of the chosen depth: the same excess over the null held at the top 100, 500, and 1,000 ranked genes per method (**Supplementary Fig. 1d**). Pairwise overlaps between datasets likewise exceeded the expectation for every dataset-pair (**Supplementary Fig. 1f**). Genes that exhibit clonal memory are therefore not cell type-specific, although the overlap remains partial, as expected for gene sets called in different tissues and organisms.

To understand what distinguishes memory genes, we first analyzed expression levels. Consensus memory genes were more highly expressed than the non-memory genes in all four datasets (**Fig. 1e**; two-sided Mann-Whitney U, p < 0.001). We therefore matched on expression throughout the remainder of this work, comparing memory genes with non-memory genes drawn from the same expression distribution. Against such a background, memory genes were expressed in fewer cells than their matched controls in all four datasets, whereas protein abundance^35^ and exonic gene length did not differ consistently (**Supplementary Fig. 1g**).

Subsequently, we compared consensus memory and expression-matched non-memory genes across methods on molecular properties, using published genome-wide measurements of mRNA half-life^36^, promoter CpG content^37^ and transcriptional bursting^38^ (**Fig. 1f**). mRNA transcripts of memory genes were longer-lived in all but one dataset group (p < 0.05 to p < 0.001; empirical two-sided permutation p-values). Transcriptional burst frequency was unchanged (p ≥ 0.05) or lower in memory genes (p < 0.05), and burst length did not differ in any group (p ≥ 0.05), so memory thus does not seem to be maintained by more frequent or longer transcriptional bursts. The promoters of memory genes, by contrast, were significantly CpG-depleted in every group (all p < 0.001). We additionally tested 34 sequence, annotation and gene-architecture features in the same expression-matched comparison (**Supplementary Data S1**; **Methods**). Promoter CpG depletion was the largest and most consistent difference, with all eight CpG measures lower in memory genes in every dataset. Memory genes also had fewer isoforms (all four datasets) and shorter gene lengths (three datasets). Two of these differences could explain how expression states persist: stability captures the turnover of a gene’s expression state, while CpG content may encode that state in DNA methylation, which is heritable across divisions^15^; neither reflects a higher transcription rate, consistent with unchanged bursting.

We next asked what memory genes encode, testing each dataset’s consensus memory gene set for overrepresentation of Gene Ontology^39,40^ terms against that dataset’s own measured transcriptome (one-sided Fisher’s exact test, Benjamini-Hochberg across terms; mouse sets tested in the mouse ontology; **Methods**). In every dataset, memory genes were enriched for tissue-related terms: myeloid cell differentiation in T-cell progenitors (732 terms at q < 0.05), cytoplasmic translation and muscle cell differentiation in melanoma (942), leukocyte migration in HSPCs (850), and gliogenesis in brain (627). Across datasets, memory gene ontology enrichments converged: 94 of the 4,278 Biological Process^39,40^ terms measurable in every dataset were significant in all four, and 291 in at least three. In the smaller Cellular Component ontology^39,40^, eight out of 650 terms were significant in all four datasets, six of them related to the endomembrane system, and the remaining two to the cell surface and extracellular matrix (**Supplementary Data S2**).

To determine how many generations the transcriptional memory survives, we measured the expression autocorrelation as the intraclass correlation coefficient among cells of the same clone and stratified clones by size (**Fig. 1g**). Because our data are snapshots, the number of divisions separating cells of a clone cannot be observed directly and has to be estimated from clone size: we assume that measured clone sizes are proportional to true clone sizes, and a clone of *n* measured cells has undergone at least log_2_(*n*) divisions. We therefore binned clones by size and placed each bin on a generation axis at its mean log_2_ clone size. Memory genes retained higher autocorrelation than non-memory genes across several divisions in all four datasets, and their autocorrelation tended to decay more slowly.

The features that distinguish memory genes from others are intrinsic to the transcript and its promoter. We next asked whether memory genes have a special regulatory role. Single-cell pooled CRISPR screens measure the transcriptome-wide effect of thousands of single-gene knockdowns or knockouts, allowing two distinct questions: (i) whether perturbing memory genes produces different effects from perturbing other genes, and (ii) how memory genes themselves respond when other genes are perturbed. To answer these questions, we paired each lineage dataset with at least one Perturb-seq screen in the same cell type, giving six screens and 9,551 CRISPR knockout or CRISPRi knockdown perturbations across T cells^28,30^, melanoma^31,32^,

HSPCs^33^, and brain^29^ (**Fig. 2a, Supplementary Table S3**). We first analyzed if perturbing memory genes generated a different response as compared to perturbing non-memory genes (**Fig. 2b**), scoring each perturbation by the mean absolute log fold change (LFC) it produced across all measured genes. Perturbing a memory gene produced a transcriptome-wide response that was no different than perturbing any other gene, and if anything smaller: the median impact differed by-5% in T cells (Zhu^28^),-8% in melanoma (Frangieh^32^),-3% in HSPCs^33^, none statistically significant, and by-12% in brain^29^, the only significant difference (**Fig. 2c**). The same held true when memory genes were defined within each experimental group instead of being merged across groups, across nine screen-by-group comparisons (**Supplementary Fig. 2a**). The two remaining screens^30,31^ contained only one and five memory-gene perturbations, respectively; they are shown but were not statistically tested (**Fig. 2c**). Memory genes are therefore not distinguished by the size of the response they cause.

**Fig. 2:**
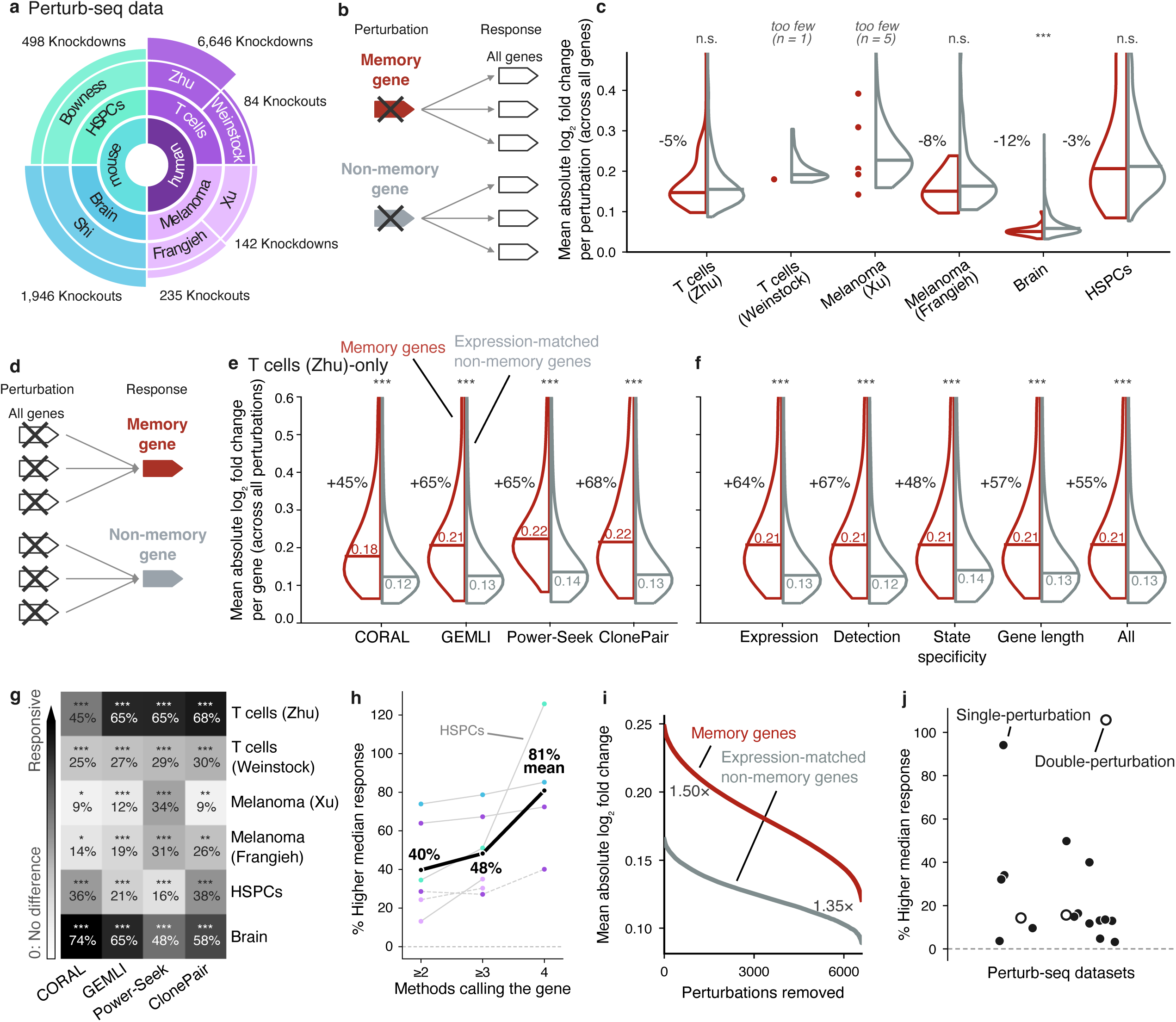
Memory genes respond more strongly to perturbations than matched control genes, whereas perturbing them does not induce larger transcriptome-wide responses. **a** Perturb-seq screens analysed, at least one per tissue, paired with the lineage datasets from Fig. 1b: 9,551 perturbations in six screens (Zhu^27^, Shi^28^, Bowness^29^, Frangieh^32^, Xu^31^, Weinstock^30^). Rings from the centre: species, tissue and screen (first author); radial extent of each screen segment scales with the logarithm of its number of perturbations, given alongside. Perturbations are CRISPR knockouts (KO) or CRISPRi knockdowns (KD), as indicated. **b** Schematic of the comparison in **c**: each perturbation is scored by the transcriptome-wide response it produces, and perturbations of memory genes (red) are compared with all other perturbations in the same library (grey). **c** Transcriptome-wide impact of each perturbation, one split violin per screen. Red, perturbations of consensus memory genes, defined as genes called by at least two of the four methods in the lineage dataset matched to that screen; grey, perturbations of genes called by no method. Impact is the mean absolute log_2_ fold change a perturbation produces across all measured genes. Horizontal lines, medians; percentages, the difference in median relative to the grey set; asterisks, two-sided Mann-Whitney U test with Benjamini-Hochberg correction across screens (n.s., q > 0.05; *, q < 0.05; **, q < 0.01; ***, q < 0.001). Screens with fewer than ten memory-gene perturbations are shown but not tested: their perturbations are drawn as individual points against the same grey distribution, with the count given in place of the asterisks (T cells (Weinstock), n = 1; melanoma (Xu), n = 5). **d** Schematic of the comparisons in **e**-**j**: each gene is scored by how strongly it responds across all perturbations of a screen, and memory genes (red) are compared with control genes matched on expression (grey). **e** Response of memory genes and expression-matched control genes in the T cell screen (Zhu), for each method separately. Response is the mean absolute log_2_ fold change of a gene across all single-gene perturbations of the screen. Memory genes are the top 300 of each method in the matched lineage dataset (T cells, Michaels^22^); controls are drawn from the genes no method placed in its top 300, matched to the memory set’s expression distribution by quantile binning (50 bins, averaged over 100 draws; **Methods**). Red, memory genes; grey, matched controls; horizontal lines, medians; percentages, the increase in median over the matched control; asterisks, two-sided Mann-Whitney U test with Benjamini-Hochberg correction across the panel (***, q < 0.001). The y-axis is shared with **f** and truncated at 0.6 for display; medians, percentages and tests are computed on the complete distributions. **f** Response of consensus memory genes, genes called by at least two of the four methods, against control sets matched on different gene properties; same screen, axis and conventions as in **e**. Controls are matched on mean expression, single-cell detection rate, cell-state specificity, gene length, or all four jointly and are drawn by nearest-neighbor matching (k=10) among genes called by no method. **g** Increase in median response of memory genes over expression-matched controls, for every method (columns; top 300 genes per method) and every lineage dataset paired with a screen from the same tissue (rows). Rows are named by tissue with the screen’s first author; colour and number, the percentage increase, with zero marking no difference; asterisks, two-sided Mann-Whitney U test with Benjamini-Hochberg correction (*, q < 0.05; **, q < 0.01; ***, q < 0.001). **h** Increase in median response of memory genes over expression-matched controls, with memory genes defined by increasing method agreement: genes ranked among the top 300 by at least two, at least three, or all four methods. One line per lineage-screen pairing, that is one row of **g**; black, the mean across pairings. The four-method mean covers four of the six pairings, the other two having fewer than ten genes at that level. **i** Response of memory genes and expression-matched controls as perturbations are removed in order of their impact on memory genes, strongest first. The 6,646 perturbations of the T cell screen (Zhu) were ranked by their mean absolute log_2_ fold change across consensus memory genes; at each point on the x-axis that many top-ranked perturbations have been removed. Red, mean absolute log_2_ fold change of memory genes over the remaining perturbations; grey, the same for an expression-matched set of genes called by no method (50 expression bins). Ratios between the two curves are given at the start and end. Means over fewer than 50 remaining perturbations are unstable and are not shown. **j** Response of memory genes identified without lineage information, in 15 Perturb-seq screens (18 comparisons) comprising 68,659 perturbations and 13.3 million cells. Power-Seek²¹ was run on each screen’s own unperturbed cells and its top 300 genes compared with an equal-sized, expression-matched set of genes it did not call. Response is the mean absolute log_2_ fold change of a gene across all perturbations of that screen, with each perturbation’s own target masked, over genes above 0.1 UMI in the control cells. Each point is one comparison. Filled, a screen scored on perturbations that each target one gene. Three of the screens also contain perturbations that target two genes at once; each of these contributes a second, open point scored on its two-gene perturbations only, which is why 15 screens give 18 comparisons. The y-axis gives the percentage by which the memory-gene median exceeds that of the matched background; dashed line, no difference. Two-sided Mann-Whitney U test with Benjamini-Hochberg correction across screens; **Supplementary Fig. 7** names each dataset and gives its medians, effect sizes and per-screen statistics.

To test how strongly memory genes respond to perturbations of other genes, we scored every gene by its mean absolute LFC across all single-gene perturbations of a screen, masking each gene’s own perturbation where applicable (**Fig. 2d**). In the best-powered T cell screen with 6,646 perturbations, the top 300 memory genes had significantly higher median responses than their expression-matched controls for every method: +45% (CORAL^20^), +65% (GEMLI^21^), +65% (Power-Seek^22^) and +68% (ClonePair) (**Fig. 2e**; all p < 0.001, two-sided Mann-Whitney U test). The paragraphs that follow test this result against potential confounders, further screens and tissues, and alternative ways of identifying memory genes.

To test confounding explanations, we additionally matched the control set on single-cell detection rate, cell-state specificity, gene length, or all properties jointly instead of expression alone. The effect persisted at +48% to +67% for the 336 consensus memory genes (**Fig. 2f**). Repeating it with consensus memory genes called within each experimental group separately gave +39% to +62% (**Supplementary Fig. 2b**), and on the memory set returned by each method separately, +31% to +71% across all twenty combinations of the four methods and five confounder matching schemes (**Supplementary Fig. 2c**). Expression-matched controls remained available across the whole expression range, even above the 95th percentile of memory gene expression (**Supplementary Fig. 3a**). The effect also held across other analytical choices: across expression floors from 0 to 2 UMI per control cell, memory genes exceeded their controls by 38% to 64%, and across alternative summaries of a gene’s response, including the median, the standard deviation and the maximum absolute log_2_ fold change, by 41% to 300% (**Supplementary Fig. 3b**). Computing response separately within each of the four donors of the T cell screen gave on average +35% to +66% across methods (**Supplementary Fig. 3c**). Across these comparisons, memory genes exceeded their matched controls by at least 31%. Their stronger response therefore does not depend on how controls are matched, how memory genes are called, or how a gene’s response is quantified.

We next checked whether the effect generalized across tissues and screens. In all 24 combinations of four lineage datasets and six matched screens, and four memory gene calling methods, memory genes were significantly more responsive than their expression-matched controls, by +9% to +74% (**Fig. 2g**). The result held when memory genes were called within each experimental group separately, for 46 of 52 method-by-group comparisons (**Supplementary Fig. 4a**), and at every ranking depth from top 100 to top 2,000 memory genes per method (**Supplementary Fig. 4b**).

To test whether the effect grew with increasing agreement between methods, we averaged across the six lineage-screen pairings. The excess median response over matched controls rose from +40% for genes called by at least two methods to +48% at three and +81% at all four (**Fig. 2h**). The same trend appeared when memory genes were defined within individual experimental groups, where the average rose from +36% at two methods to +46% at three and +65% at four (**Supplementary Fig. 4c**).

To test whether a few perturbations drove the effect, we ranked the 6,646 perturbations of the T cell screen by their impact on consensus memory genes and removed them successively, strongest first. The higher response of memory genes over their expression-matched controls, at 50% with all perturbations included, remained above 35% with perturbations removed (**Fig. 2i**), and between 22% and 37% when each method was considered separately (**Supplementary Fig. 4d**).

As an orthogonal measurement, we repeated the perturbation response comparison against expression-matched controls with memory genes that were determined experimentally with complementary approaches. In an independent, previously published experiment, MemorySeq^16^ identified 227 memory genes in melanoma cells. These genes significantly overlapped with the memory genes we identified computationally in the lineage-barcoded melanoma dataset, compared to an expression-matched baseline (**Supplementary Fig. 5a**). These genes were themselves more responsive than their matched controls in both melanoma screens, by +28% (Xu) and +36% (Frangieh), compared to +11% and +23% in the lineage-barcoded set (**Supplementary Fig. 5b**).

To test generality further, we removed the lineage-barcode requirement altogether and called memory genes directly in the unperturbed control cells of each perturbation screen. With no tissue-matched clonal dataset needed, this opened 15 additional screens to analysis. We used three methods that require no lineage barcodes: Power-Seek, GEMLI’s lineage-free mode, and isCORAL, CORAL’s lineage-free update, which infers pseudoclones^41^. In four out of the five screens with a matched lineage dataset, the resulting memory gene sets were at least as responsive as their barcode-derived counterparts (**Supplementary Fig. 6a**). Genes called by all three in control cells exceeded their matched controls by +127% (T cells), +136% (melanoma, Frangieh), +68% (melanoma, Xu), +148% (HSPCs) and +212% (brain) (**Supplementary Fig. 6b**). Across the additional 15 screens, comprising 68,659 perturbations and 13.3 million cells, memory genes were more responsive than expression-matched controls in every screen tested, by +3% to +94% for single-gene perturbations and by +14% to +106% when scored on paired perturbations alone (**Fig. 2j, Supplementary Fig. 7, Supplementary Data S5**). Together, across every dataset, method, matching scheme and screen we tested, memory genes responded more strongly to perturbation than comparable genes.

To understand why memory genes respond more strongly to perturbations, we devised a mathematical response model and validated it across perturbation screens (**SI**). A perturbation screen yields a matrix of gene expression log_2_ fold changes (LFCs), with one row per perturbation and one column per measured gene. We summarized the response of a gene set G to a perturbation *i* by the root-mean-square (RMS) of its entries in that row. Although individual genes may respond differently, we hypothesized that this quantity approximately factorizes as R*ᵢ*(G) ≈ G*ᵢ*H_G_, where G*ᵢ* describes potency of perturbation response and H_G_ describes gene-set susceptibility (**Supplementary Fig. 8a**). If responses separate in this way, the ratio of memory-gene to non-memory-gene responses is independent of perturbation identity, as the potency factor cancels out. Henceforth, subscripts denote memory (mem) and non-memory (nonmem) gene sets. By taking ratios of RMS responses, we can estimate G_mem_/G_nonmem_ and H_mem_/H_nonmem_, as well as a factorization constant C predicted to equal unity (**SI**). Across all 23 datasets with perturbed memory genes, the factorization constant was close to one (C = 0.98 ± 0.01, mean ± SEM), consistent with approximately separable responses. Memory-gene perturbations were modestly less potent than non-memory perturbations (G_mem_/G_nonmem_ = 0.92 ± 0.02, p = 0.0005), whereas memory genes were substantially more susceptible as targets (H_mem_/H_nonmem_ = 1.51 ± 0.07, p < 0.0001; **Supplementary Fig. 8b**).

We next asked why responses separated in this way. To determine whether different perturbations produce a common pattern of gene expression changes, we used singular-value decomposition to separate the overall magnitude of each response from its shape. We applied this analysis at two scales, using 1) signed LFCs, and 2) gene populations (unsigned RMS responses for memory genes and five non-memory quintiles ordered by mRNA half-life or CpG content). At the LFC level, the leading singular mode explained only 9.1% of the variation in non-memory genes and 12.6% in memory genes, indicating that the identities and directions of responding genes differ substantially across perturbations. At the population level, however, the leading mode explained 99.7% of the variation (**Supplementary Fig. 8c**). Thus, individual genes may respond differently to each perturbation, but the relative distribution of response magnitudes across gene populations remains remarkably consistent. Consistent with this separability, susceptibility ratios that were fit on training perturbations accurately predicted held-out RMS responses across 24 datasets (median R^2^ = 0.96 in log space, 454,150 predictions; **Supplementary Fig. 8d**). Nevertheless, perturbations produced distinct but correlated gene-level response patterns (**Supplementary Fig. 8e**), suggesting that collective structure may underlie the enhanced susceptibility of memory genes. We reasoned that memory genes might respond more strongly either as independent entities or through a shared regulatory component. These possibilities can be distinguished by the growth of the leading eigenvalue of their response correlation matrix as genes are added: independent responses yield little growth, whereas a shared component produces approximately linear growth. In K562 CRISPRa, this growth was 3.4-fold steeper for memory genes than matched controls (**Supplementary Fig. 8f**). Across 24 datasets, memory genes exhibited a 2.1-fold greater mean spectral scaling slope (0.31 ± 0.09 versus 0.15 ± 0.03, mean ± SEM; p = 0.021, paired t-test; **Supplementary Fig. 8g**). Together, these results suggest that memory genes respond more strongly because they exhibit enhanced collective susceptibility, rather than because perturbations targeting them are intrinsically more potent. A slowly relaxing regulatory component could therefore both preserve gene-expression states and amplify their response to perturbation, linking cellular memory and responsiveness through a common mechanism.

## Discussion

Here we set out to ask whether methods that call clonally heritable or memory genes by independent statistics identify the same genes, what distinguishes those genes, and how they act within regulatory networks. The methods converged on a common set in every dataset, and some of those genes recurred across tissues and species. What distinguished memory genes from non-memory genes was not how their expression was produced but how slowly it turned over: longer mRNA half-lives and CpG-depleted promoters, with burst length unchanged and burst frequency unchanged or lower. Perturbing memory genes produced no larger response than perturbing any other gene, but when other genes were perturbed, memory genes responded more strongly than matched controls in every screen, memory-gene calling method, and matching scheme we tested.

Stability across divisions suggests that memory genes should resist perturbation, acting as “pillars” that hold cell identity by moving less than other genes. Instead, we found that they act as “springs”, displaced further than other genes. Stability and displacement may follow from the same timescale: a gene accumulates a sustained input for as long as it remembers, so the genes that remember longest are displaced furthest.

Memory genes appear to be receivers, not drivers: they report cell state but are not obvious targets for changing it, though their expression in fewer cells may dilute the measured effect of perturbing them. The distinction matters wherever cells are deliberately moved. Heritable expression states have already been linked to therapy resistance^5–9^, to which cells reprogram to pluripotency^10,11^, and to susceptibility to viral infection^12,13^. Each of these follows a sustained, directed input, and memory genes should move furthest under such inputs. That makes them candidate reporters for how a population is responding, whether to a reprogramming protocol, a drug or an infection. Memory is also visible in perturbation data alone: genes called in the unperturbed control cells of a screen, with no lineage information, responded as strongly as those called from barcoded clones, so memory genes can be identified without a clonal experiment.

Current methods resolve clone membership but not the relationships within a clone, so sisters and distant relatives contribute equally. Phylogenetically resolved lineage tracing would give the decay of memory with division number directly, rather than inferring it from clone size, and is the most direct route to the timescale we did not measure. In tissues where cells move little, spatial proximity stands in for lineage proximity, though neighboring cells also share a niche. Spatial CRISPR screens^42–44^ would then allow memory genes to be called and perturbed in the same tissue. Memory genes are defined here on transcripts, while the CpG depletion we observe implicates the upstream process of DNA methylation, which we did not measure. Methylation at individual CpGs is clonally heritable and can now be read alongside expression in the same cells^45^, so measuring both in lineage data and in perturbation screens alike^46^ would show which layer holds the state and which moves when the cell is perturbed. In our study, promoter CpG content was computed directly from the primary assembly and was therefore available for every gene, whereas the mRNA half-life and bursting estimates were drawn from measurements in other cell types and covered a subset of genes. This limits what those comparisons can support.

## DATA AND CODE AVAILABILITY

All code needed to reproduce the analyses and figures in this study is available at https://github.com/marrlab/PERM. All preprocessed datasets and supplementary data are publicly available under 10.5281/zenodo.24590499 and are summarized in **Tables S1, S3, and S4**.

## AUTHOR CONTRIBUTIONS

IB, CM and YG conceived and designed this study. IB and VN performed the analysis with input from CM and YG. BKS formulated the analytical model with input from IB, CM and YG. IB, VN, CM and YG wrote the manuscript and prepared the figures with input from BKS.

## ACKNOWLEDGEMENTS

We thank Magdalena Götz (Munich), Madeline Melzer (Chicago), and members of the Goyal, Marr and Colomé-Tatché labs for helpful discussions and comments on the manuscript. We thank the Feinberg Information Technology, Northwestern University Information Technology, and Quest High Performance Computing Cluster at Northwestern University Feinberg School of Medicine for computational support and services. YG thanks the National Science Foundation through the Center for Living Systems Ignite Award (grant no. 2317138) as well. This work was supported by the National Institute for Theory and Mathematics in Biology (NITMB) through the National Science Foundation (NSF; DMS-2235451) and the Simons Foundation (MPTMPS-00005320). BKS was supported by the Northwestern University Clinical and Translational Sciences Institute (NUCATS) T32 program, NITMB, Biohub SPARK award, and Northwestern University startup funds to YG. YG acknowledges grant support from the Pew-Stewart Scholars Program, American Heart Association, Biohub Investigator Award, Edward Mallinckrodt, Jr. Foundation Grant, and Burroughs Wellcome Fund Career Awards at the Scientific Interface. CM received funding from the European Research Council under the European Union’s Horizon 2020 Research and Innovation Programme (grant agreements 866411, 101113551, and 101213822) and support from the Hightech Agenda Bayern. This work was supported through the Bavarian AI Foundation Model Initiative, funded by the Bavarian State Ministry of Science and the Arts (StMWK). IB is supported by the Helmholtz Association under the joint research school “Munich School for Data Science - MUDS” and thanks the Biomedical Center at Ludwig-Maximilians-Universität for computational services and support.

## AI USE

During the preparation of this work, the authors used Anthropic’s Opus 4.8, 5, and 5.5 and Sonnet 4.6 via Claude Code to write and edit code, and via Claude Chat to brainstorm, search the literature, and proofread the manuscript. The authors reviewed and edited AI-generated content as needed and take full responsibility for the content of the published article.

## Supplementary Data

- Suppl. Data S1. Gene features

- Suppl. Data S2. GO analysis tables

- Suppl. Data S3. Lineage-barcode datasets table

- Suppl. Data S4. Main Perturb-seq screens table

- Suppl. Data S5. Perturb-seq screens used in generalization analysis table

- Suppl. Data S6. Memory gene lists per method, dataset and group

- Suppl. Data S7. Consensus memory gene sets

- Supporting Information (SI). Mathematical response model

## Methods

### Lineage-barcoded scRNA-seq datasets

We assembled four published lineage-barcoded scRNA-seq datasets across two organisms (human and mouse), four barcoding systems (LARRY^24^, STICR^25^, the transcribed barcode^23^, and transcribed semi-random barcodes^17^), and both developmental and cancer settings (**Fig. 1b** and **Table S1**).

#### Human haematopoietic and T-cell progenitors, transcribed barcodes

Michaels et al. differentiated human induced pluripotent stem cells towards blood and progenitor T-cell fates, using transcribed lineage barcoding to link clonal relationships to transcriptional states during fate restriction^23^. At day 0, the authors seeded 7,500 CD34^+^ cells, derived from the iPS11 induced pluripotent stem cell line, into one well and labelled them with the lineage barcode library on day 3. Two days after transduction (day 5), half of the well was sampled for scRNA-seq (time point 1). On day 7, the remaining cells were transferred to DLL4-and VCAM1-coated plates to begin progenitor T-cell differentiation. Three days later (day 10), a subsample of 16,500 cells was taken for scRNA-seq (time point 2). Both samples were profiled with the 10x Chromium Single Cell 3’ v3.1 chemistry, and lineage barcodes were additionally recovered by targeted amplification. After quality control, the two time points comprised 3,948 barcoded singlets and 23,163 genes. Clone sizes were computed within each time point, yielding 1,040 clones at timepoint 1 (836 clones with >= 2 cells) and 620 clones at timepoint 2 (451 clones with >= 2 cells).

#### Human melanoma, scMemorySeq

Harmange et al. coupled lineage barcoding to scRNA-seq to quantify how single-cell expression states persist and to identify cells that switch state^17^. They sorted two starting populations of 1,000 barcoded WM989 melanoma cells each: one enriched for the rare primed state (the top 0.2% of EGFR-and NGFR-expressing cells, labelled E in the authors’ code) and one mixed population of all live cells (M), consisting predominantly of drug-susceptible cells. Both were expanded for four to five divisions before capture. Expression counts from the resulting 12,531 drug-naive WM989 cells and 33,538 genes were downloaded from GEO (GSE237228, GSM7597476). The lineage-barcode counts are not deposited there and were taken from the Seurat object released with the authors’ analysis code (see code availability of corresponding paper^17^). Cells were assigned to clones using the authors’ assignment code, with a single change: the minimum clone size was set to 2 rather than 4, so that two-and three-cell clones were retained. Run with the authors’ original threshold, our pipeline reproduced their published assignments almost exactly (adjusted Rand index^47^ 0.9993). This yielded 8,000 cells in 450 clones, with each clone confined to either the E or the M population. We therefore analyzed the E (2,226 cells) and M (5,774 cells) populations separately.

#### Mouse hematopoietic progenitors, LARRY

Weinreb et al. tagged Lin^-^Kit^+^Sca1^+^ (LSK) hematopoietic stem and progenitor cells with an expressed lentiviral barcode library and profiled them by inDrops across an in vitro differentiation time course, linking clonal identity to fate outcome^24^. Because the barcodes are transcribed, they are recovered from the same library as the transcriptome rather than by separate amplification. We used the three barcoded collection points of the culture (days 2, 4, and 6; 15 sequencing libraries) together with the authors’ published clone assignments, and removed cells carrying no assigned barcode. This left 24,908 barcoded cells and 25,289 genes (GEO, GSE140802), carrying 3,720 distinct barcodes, forming 1,625 clones at day 2 (863 of at least two cells), 2,497 at day 4 (1,674 of at least two cells), and 2,630 at day 6 (2,150 of at least two cells).

#### Mouse forebrain, STICR

Bandler et al. tagged mitotic progenitors lining the embryonic mouse ventricle with the STICR barcode library and profiled their postnatal progeny, allowing clonal relationships to be traced across the major forebrain cell classes^25^. The published data combine two barcoding assays, STICR and TrackerSeq. We used the STICR libraries only: TrackerSeq contributes a single library of 1,798 cells, labelled at E12 and collected embryonically at E16, whereas every STICR library was collected postnatally. We rebuilt an AnnData object from the 22 barcoded libraries of GSE188528 (six wild-type regional dissections without barcodes were excluded), yielding 26,124 cells and 17,702 genes, of which 21 libraries and 24,326 cells are STICR.

Because clones are nested within labeling age, labeling age defines the groups on which memory calling methods are run. We therefore restricted the analysis to a covariate-matched core in which collection age (P7), strain (SW), and dissection (whole forebrain) are held constant, so that labeling age is the principal variable distinguishing the groups. Embryonic days 13 and 14 are the only labeling ages for which such a matched comparison exists: E10 is entirely C57BL/6 collected at three collection ages, while E12 spans five libraries, five collection ages and two strains, and is also the only age. Without this restriction, the embryonic day 13 group would span collection ages P5-P14, and the embryonic day 14 group would mix whole-forebrain with region-dissected libraries, which contribute 43% of its sister pairs. The matched core comprises 10,170 cells in eight libraries: embryonic day 13 (2,147 cells; 335 clones of >= 2 cells) and embryonic day 14 (8,023 cells; 1,434 clones of >= 2 cells).

**Table S1:** Lineage-barcoded scRNA-seq datasets. The four lineage-barcoded datasets on which the memory-gene callers were run. Groups are the experimental units each dataset is divided into, and every quantity is computed within a group. A clone observed at two timepoints counts as two clones, because the callers compare cells within a group and never pool across groups. Cells are those carrying an assigned barcode after quality control. Clones give all clones, with those of at least two cells in brackets, since one-cell clones carry no within-clone information and are used by none of the methods. The last column names the Perturb-seq screen each dataset was paired with for the response analysis, matched by cell type (**Table S3**). Full dataset metadata, including repositories and file-level details, are in **Supplementary Data S3**.

| **Dataset** | **Publication** | **Accession** | **Barcoding assay** | **Species, system** | **Groups** | **Cells** | **Clones (≥2 cells)** | **Cells in clones of ≥2** | **Median clone size** | **Paired Perturb-seq dataset** |
|---|---|---|---|---|---|---|---|---|---|---|
| Michaels | Michaels et al. 2024, *Cell Systems* 15(12):1245-1263 | GSE248549 | transcribed lineage barcoding | human, hPSC-derived haematopoiesis | Timepoint 1 and 2 | 3,948 | 1,660 (1,287) | 3,575 | 2 | Zhu, Weinstock |
| Harmange | Harmange et al. 2023, *Nature Communications* 14:7130 | GSE237228 | scMemorySeq | human, WM989 melanoma | enriched, mixed | 8,000 | 450 (450) | 8,000 | 9 | Xu, Frangieh |
| Weinreb | Weinreb et al. 2020, *Science* 367(6479) | GSE140802 | LARRY lentiviral barcoding | mouse, haematopoietic progenitors | days 2, 4, 6 | 24,908 | 6,752 (4,687) | 22,843 | 2 | Bowness |
| Bandler | Bandler et al. 2022, *Nature* 601(7893):404-409 | GSE188528 | STICR retroviral barcoding | mouse, forebrain | embryonic days 13 and 14 | 10,170 | 6,887 (1,769) | 5,052 | 1 | Shi |

### Preprocessing

#### Quality control and normalization

All datasets were converted to a single AnnData schema (log-normalized expression in *.X*, raw integer counts in *layers[‘counts’]*, and harmonized *clone_id*, *clone_size,* and group annotations in.*obs*). For each dataset, all methods received identical cells and identical clone labels. Quality control followed the same scheme for all four datasets: a minimum number of detected genes per cell (set per dataset, as median library sizes range from 4,424 to 54,731 UMI per cell), exclusion of cells with anomalously high gene counts as probable doublets, and a ceiling on mitochondrial content between 12.5% and 30%. Counts were normalized to 10,000 per cell and log-transformed (*normalize_total(target_sum=1e4), log1p*^48^), with raw integer counts retained in *layers[‘counts’]* for the methods that require them, and per-cell normalized totals were verified to equal 10,000 in every object. Gene selection was left to each method, since every method applies its own expression or detection criteria: a minimum mean expression (CORAL^20^), a minimum detection fraction (GEMLI^21^, ClonePair), or a minimum number of expressing cells followed by selection of the most variable genes (Power-Seek^22^). These criteria admit between 2,500 and 13,000 genes per group, so agreement between methods is bounded by the intersection of the gene sets they admit.

#### Gene identifiers

Each expression matrix was kept in the gene namespace of its species of origin, and no gene was removed on the basis of orthology. Mouse genes were mapped to human symbols only where gene lists were compared across datasets (see Comparison across methods and datasets).

#### Lineage schema

Each dataset stores, per cell, the clone assignment made by the original authors (*clone_id*, stored as a string), the clone size (*clone_size*), and an analysis group. The analysis group defines the subsets within which every method was run independently: time point for Michaels et al. (timepoints 1 and 2) and Weinreb et al. (timepoints 1, 2, and 3), labeling age for Bandler et al. (embryonic days 13 and 14) and starting population for Harmange et al. (enriched, mixed). Cells without an assigned barcode were removed before any analysis, so that only clone-assigned cells were retained. Clone size was computed within each analysis group. A clone sampled at several time points therefore counts separately at each one, and its size is not inflated by pooling.

### Identifying memory genes

We benchmarked four methods for identifying memory genes, defined as genes whose expression is inherited across cell divisions and is therefore more similar among clonally related cells than among unrelated cells. Three are published: CORAL^20^, GEMLI^21^, and Power-Seek^22^; the fourth, ClonePair, was developed by us. CORAL and ClonePair use experimentally determined clone labels, whereas Power-Seek infers memory genes from expression alone. GEMLI offers a lineage-free mode (**Supplementary Fig. 6**); however, we use GEMLI with clone labels in all main analyses. Each published method was executed as released by its authors. All methods received the same cells and clone assignments and were run separately for each experimental group in a dataset: clonally related cells tend to share a group, so pooling groups would let genes that merely distinguish cell states appear as memory genes.

We defined sister-concordant genes as genes whose expression is more similar between clonally related cells than between unrelated cells. Within each group *g* (experiment, timepoint, labeling age or starting population, as defined per dataset above), we formed, for every clone with *k* cells, all C(*k*,2) cell pairs and referred to them as sister pairs. We only consider clones with 2-8 cells. Clones sampled at two timepoints contribute pairs to each group separately. Genes detected in fewer than 5% of the cells of a group were not tested in that group. For each tested gene *i*, we computed the mean absolute expression difference across the *n* sister pairs, d*_i_*^sis^ = (1/*n*) Σ_{p=1}_^n^ |x*_i,p1_* − x*_i,p2_*|, where *n* is the number of sister pairs, *p_1_* and *p_2_* are the two sisters of pair *p*, and x*_i,p1_* is the log-normalized expression of gene *i* in cell *p_1_*. A matched null d*_i_*^rand^ was obtained from *n* random pairs of cells from the same group, drawn without replacement for groups with at least 2*n* cells, and with replacement otherwise (excluding self-pairs), so that null and observed sets always contain the same number of pairs. Restricting both sets to one group prevents between-group expression differences from inflating the null. Random pairing was repeated 100 times with a fixed seed. In each iteration, the per-pair absolute differences of sister and random pairs were compared with a one-sided two-sample t-test (sister < random), and p-values were corrected across tested genes by the Benjamini-Hochberg^49^ procedure at an FDR of 5%. For each gene, we recorded the concordance frequency π_i_, i.e., the fraction of iterations in which it was significant, and called a gene sister-concordant if π_i_ ≥ 0.80. The concordance frequency is strongly bimodal, with most genes at or close to 0 or 1 (**Supplementary Fig. 1b**), so the majority of genes are assigned identically across a wide range of thresholds. The value 0.80 was fixed in advance, and the number of genes called decreases smoothly as the threshold is raised (**Supplementary Fig. 1a**). As an effect size, we report the dissimilarity ratio r*_i_* = d*_i_*^sis^ / d*_i_*^rand^, which is below 1 for concordant genes and decreases with stronger concordance; where a ranking is required (e.g., top-N comparisons with the other methods), genes were ordered by increasing r*_i_*.

In clones of more than two cells, the same cell contributes to several pairs, so the pairs are not independent: the number of independent observations is closer to the number of clones than to the number of pairs, and the per-iteration t-test therefore returns smaller p-values than it should. We use π_i_ only as a cut-off for calling genes, and rank genes by r_i_, which does not depend on how many pairs a clone contributes. The calls are not interpreted as significance statements on individual genes: every downstream analysis compares gene sets against matched nulls, and the called sets are validated against a permutation null by their agreement with the other three methods (**Fig. 1c**).

CORAL^20^ quantifies the proportion of a gene’s expression variance that is attributable to clonal identity. From a log-normalized expression matrix and a clone label per cell, it computes for each gene the one-way ANOVA effect size w^2^ = (SS_b_ − (k−1)MS_w_)/(SS_t_+MS_w_) where *SS_b_* is the between-clone sum of squares, *SS_t_* the total sum of squares, *MS_w_* the within-clone mean square (the within-clone sum of squares divided by its *N - k* degrees of freedom), *k* the number of clones and *N* the number of cells. w^2^ estimates the fraction of total expression variance attributable to clonal identity, corrected for the variance that *k* arbitrary groups would explain by chance. Negative estimates, which arise when between-clone variation is no larger than within-clone variation, are set to zero. A null distribution is generated by permuting clone labels across cells and recomputing w^2^. Genes whose w^2^ exceeds the third quartile plus 1.5x the interquartile range of the pooled null, and whose mean expression exceeds a set minimum are reported as memory genes.

We ran *run_coral_ground_truth_analysis()* from the CORAL R package (github.com/YongjieLin1998/CORAL, commit 68b30dc) in R 4.4.3 with Seurat 5.5.1^50^, Matrix 1.7.6 and ComplexHeatmap 2.22.0^51^, with lineage barcodes as clone labels and the parameters clone_size_cutoff = 2, min_gene_mean = 0.05, permutation_repeats = 5, and the permutation seed fixed at 0. Expression was supplied in the log-normalized form described in Preprocessing, written directly into the Seurat *data* layer without further normalization. We made two modifications: (i) We removed the clone-clone energy-distance, state-clustering, and multidimensional-scaling steps, which produce CORAL’s state and MDS outputs and do not enter the memory-gene calculation. The reduced code returned per-gene statistics identical to the full version to machine precision; (ii) Clone labels were prefixed with a letter so that clones numbered 0 were not discarded as unlabelled.

Power-Seek^22^ infers memory genes from the covariance structure of expression alone, without clone labels. After each gene is standardized across cells, the cell–cell covariance matrix C = XX^T^/n_g_ is formed (X, cells x genes; n_g,_ number of genes), and each gene g is scored by how much its removal lowers the leading eigenvalues, Δ_g_ = Σ_j ≤ n_(λ_j_ − λ_j_^−g^), where *λ_1 ≥ λ_2 ≥*…are the eigenvalues of *C* in decreasing order and λ_j_^−g^ the corresponding eigenvalues of *C* recomputed with gene *g* removed. The number of eigenvalues *n* is the extent of the power-law regime of the spectrum, taken as the first maximum of the coefficient of determination (R^2^) of a log-log linear fit to the leading eigenvalues. Genes are ranked by Δ_g_.

We ran the authors’ implementation (github.com/RajuLab/CellMem-Power-Seek, commit a580d5e), with preprocessing from PowerSeekDataPrepV2.ipynb and the algorithm from PowerSeekAlgorithmOnData.ipynb. Preprocessing consisted of per-cell total-count normalization, exclusion of genes detected in fewer than 200 cells, exclusion of genes carrying any outlier in log expression under a 1.5x whisker rule applied to the 9^th^-91^st^ percentile range, and selection of the 3,000 genes with the highest coefficient of variation. The authors’ preprocessing notebook outputs the 5,000 most variable genes, whereas their analysis notebook reads a 3,000-gene matrix; we used 3,000, the input the algorithm expects. The algorithm was run with a starting eigenvalue count of 20 and proceeded in steps of 2; the 500 top-ranked genes per group were retained.

GEMLI^21^ scores memory genes using the abundance-corrected squared coefficient of variation of per-clone expression centres. For each clone within a specified size range, it takes the per-gene median across cells, computes the squared coefficient of variation of these medians across clones, regresses its logarithm on log mean expression by local polynomial regression to remove the abundance trend, and uses the residual as the gene score. An empirical p-value is obtained from random permutations of the assignment of cells to lineages. The number of permutations is fixed at 20 in the released code.

We ran *memory_gene_calling()* from the GEMLI R package (github.com/UPSUTER/GEMLI, commit d16f123; dependent only on base R and stats packages on this code path) on linear counts normalized to 10,000 per cell, with valid_lineage_sizes = 2:5 (the range the authors applied to most cell types), use_median = TRUE, and the seed fixed at 0. For all cross-method comparisons, lineage barcodes were provided rather than inferred (ground_truth = TRUE). Genes with p ≤ 0.05 were called memory genes, the criterion of the original study. Because GEMLI discards all lineages outside the specified size range, it draws on a different subset of the clonal structure than CORAL and ClonePair applied to the same data.

We additionally ran GEMLI in its lineage-free mode, which infers lineages from expression, using *predict_lineages()* (additionally requiring the packages dendextend^52^ and igraph^53^) with a sampling fraction of 0.75, 100 repetitions, a desired terminal cluster size of two to three cells, and a confidence threshold of 50, the value used in the original study. Two ambiguities in the original description were resolved as follows: The published methods specify a sampling fraction of 75%, whereas the released code defaults to two-thirds; we used 75%. The size range for scoring predicted lineages is not stated clearly, but at a confidence threshold of 50, the largest predicted lineage contained five cells, so retaining all predicted lineages and restricting to lineages of two to five cells gave identical results. Memory genes called in this mode are shown in **Supplementary Fig. 6** and are not used in any cross-method comparison.

**Table S2:** Memory gene identification methods. The four methods compared. Lineage information required states what each needs as input: CORAL and ClonePair require clone labels, GEMLI can use them or infer clones from expression alone, and Power-Seek needs none. Code gives the repository, and the commit the analysis was run from. ClonePair is introduced in this study.

| **Method** | **Publication** | **Code** | **Language** | **Lineage information required** | **What it calls a memory gene** |
|---|---|---|---|---|---|
| CORAL | Lin et al. 2026, *Cell Systems* | github.com/YongjieLin1998/CORAL, commit 68b30dc | R | yes, clone labels | genes whose expression is structurally correlated with clonal identity |
| GEMLI | Eisele et al. 2024, *Nature Communications* 15:2744 | github.com/UPSUTER/GEMLI, commit d16f123 (GEMLI_package_v0) | R | optional: clone labels, or clones predicted from expression | genes with heritable expression across a lineage (p ≤ 0.05, ranked by p and variance) |
| Power-Seek | Ghosh, Chakrabarti & Raju 2026, *Cell Systems* | github.com/RajuLab/CellMem-Power-Seek | Python | no | genes contributing to the power-law signature of the cell covariance eigenspectrum |
| ClonePair | this study | github.com/marrlab/PERM | Python | yes, sister pairs and small clones (2 to 8 cells) | genes concordant between sisters above chance, called at a concordance frequency of 0.80 over randomizations (one-sided two-sample t-test per iteration) |

### Comparison across methods and datasets

The four methods report different statistics that cannot be compared directly, so we compared the gene lists they return instead. Power-Seek^22^ returns its memory genes ranked, most confident first, by the eigenvalue contribution Δ_g_. CORAL^20^ returns an unranked set of memory genes with the effect size w^2^ of each gene, which we ranked in decreasing order of w^2^. GEMLI^21^ returns genes ordered by empirical p-value and then by its variability score. Because that p-value is computed over 20 permutations, it takes only the values 0 and 0.05 and ties thousands of genes, so we ranked by the variability score alone. For ClonePair, memory genes were ranked by increasing dissimilarity ratio *r*.

Every method returns one ranked list per experimental group, and these lists were combined into consensus lists in one of two modes, stated with each analysis. In the per-group mode (**Fig. 1c, 1f, 1g**), each group is analyzed on its own, so that a gene called in one group is not credited in another: each method’s list for that group is truncated to its top N genes, and each gene is assigned the number of methods whose truncated list contains it. In the merged mode, used where a single memory gene set per dataset is required **(Fig. 1d, 1e** and all perturbation analyses), each method’s per-group lists are first merged into one ranking in which each gene keeps its best (minimum) within-group rank; these merged lists are then truncated to their top N and combined across methods in the same way. In both modes, consensus lists contain the genes called by at least two, at least three, or all four methods. Non-memory genes are genes that occur in none of the four methods’ top N, where N is the cutoff used for the memory genes in that analysis.

Truncation is needed because list lengths differ between methods, so agreement counts would otherwise reflect list length: Power-Seek returns exactly 500 genes per group, the other three methods between 143 (ClonePair, Weinreb, timepoint 1) and 9,250 genes (CORAL, Harmange, mixed). Every method is therefore truncated to the same cutoff, N = 300 unless stated otherwise. A method that returns fewer than N genes for a group contributes all of them. This affects only one of the 36 method-by-group lists (ClonePair, Weinreb timepoint 1, 143 genes), and no list is shorter than 100 genes. In the permutation null for method agreement, each method draws as many genes as it contributed, so the shorter list is accounted for when agreement is compared by chance.

Mouse genes were mapped to human orthologs with NCBI’s curated ortholog table (gene_orthologs, downloaded 15 September 2026; 17,092 one-to-one human–mouse pairs), resolving outdated dataset symbols through NCBI gene synonyms (Homo_sapiens.gene_info and Mus_musculus.gene_info, downloaded 15 September 2026)^34^. Genes absent from the NCBI table were assigned from a HomoloGene-based map retrieved through MyGene.info^54–56^ (908 genes for Weinreb, 358 for Bandler). The map file records the source of each assignment. All human symbols, in the human datasets and after mapping, were normalized to the current NCBI symbol, converting only unambiguous synonyms. In total, 67% of Weinreb’s and 84% of Bandler’s measured genes were mapped.

### Characterization of memory genes

#### Mean expression

Consensus memory genes (merged mode, called by at least two methods; see above) were compared with non-memory genes by their mean log-normalized expression, averaged over all cells of the dataset, using a two-sided Mann–Whitney U test for each dataset.

#### Molecular features

mRNA half-lives were taken from the consensus estimates of Agarwal and Kelley^36^, who derived a cell-type-agnostic half-life per gene as the first principal component across 39 human and 27 mouse transcriptome-wide decay datasets; the human and mouse tables were matched to the human and mouse lineage datasets, respectively, by gene symbol, so no orthology mapping was needed. Transcriptional burst frequency and burst length were taken from Boe et al.^38^, which were measured in mouse; for the human lineage datasets, these values were transferred via one-to-one orthologs (see above). CpG depletion was quantified as the CpG observed/expected ratio in a promoter window centered on the transcription start site (TSS ± 500 bp), computed as (n_CpG_ x L) / (n_C_ x n_G_), where n_CpG_ is the number of CpG dinucleotides in the window, n_C_ and n_G_ are the numbers of C and G nucleotides, and L is the window length^37^. Lower ratios indicate stronger depletion. Promoter sequence was taken from the GENCODE primary assemblies (human release 47, GRCh38; mouse release M36, GRCm39) and uppercased before counting, because these assemblies are soft-masked for repeats. The transcription start site is the 5’ end of each gene’s representative transcript, rather than the gene’s most 5’ annotated start site.

Analyses were run separately for each group within a dataset (per-group mode, called by at least two methods; see above). For each feature, genes without a value were excluded, and the test statistic was the difference in mean feature value between memory and non-memory genes. To control for expression, genes were binned into 10 equal-frequency bins of mean expression and memory labels were permuted among genes within each bin, preserving the number of memory genes per bin, 10,000 times. The two-sided empirical p-value is (1 + number of permutations with |Δ| at least as large as observed) / (1 + 10,000). P-values are reported per feature and group and are not corrected for multiple testing.

#### Feature screen

To test memory genes against a broader set of gene properties (**Supplementary Data S1**), we compared, in each dataset, the consensus memory genes (called by at least two of the four methods within their top 300, merged across groups) with non-memory genes (in none of the four methods’ top 300; see above), on 34 features. CpG and promoter: overlap of the promoter with a CpG island (UCSC cpgIslandExt; TSS ± 500 bp, ± 1 kb and ± 2 kb), the CpG observed/expected ratio, CpG percentage and GC percentage of the island, distance from the TSS to the nearest island, and promoter CpG observed/expected ratio, CpG density and GC content (TSS ± 500 bp and ± 1 kb, computed as above). Annotation and constraint: publications (NCBI gene2pubmed), Gene Ontology terms in total, per aspect and excluding electronic annotations (GOA for human, MGI for mouse), protein-interaction partners (STRING v12, combined score ≥ 400 and ≥ 700), InterPro domains (reviewed UniProt entries), and LOEUF, pLI and missense Z (gnomAD v4.1; human only). Gene architecture (GENCODE, representative transcript as above): exonic length, exon and intron number, total intron length, gene and transcript span, number of annotated transcripts, protein-coding biotype, and the fraction of the gene body and of the promoter (TSS ± 1 kb) covered by phastCons elements (100-way for human, 35-way for mouse). Count features were log-transformed (log10(x + 1)). Each memory gene was matched to its ten nearest background genes in log mean expression, and the effect size was the standardized mean difference between memory genes and their matched controls (percentage points for binary features), with a 95% confidence interval from 400 bootstrap resamples of the memory genes. Significance was assessed by permuting memory labels within 50 equal-frequency expression strata (1,000 permutations); because this null reproduces the memory genes’ expression profile and is therefore not centered on zero, the two-sided p-value was computed from the observed difference’s z-score relative to the null distribution. P-values were corrected with the Benjamini-Hochberg procedure across all features within each dataset.

#### Other gene properties

Detection rate is the fraction of cells in which a gene has non-zero counts, computed over the same cells as its mean expression. Protein abundance was taken from the PaxDb v5.0^35^ *Homo sapiens* whole-organism integrated dataset, in parts per million. Ensembl protein identifiers were mapped to gene symbols with MyGene.info and isoforms collapsed by their maximum, leaving values for 18,174 genes. PaxDb provides human values only, so mouse datasets were mapped to human symbols by orthology before joining. Gene length is the exonic length of a representative transcript, preferring the MANE Select transcript^57^ where one exists and otherwise the transcript with the most exons. MANE is defined for humans only, so every mouse gene falls to the second rule. For each of these three properties, consensus memory genes (merged mode, at least two methods) were compared with non-memory genes matched one-to-one on log mean expression by nearest neighbor (k=10, each background gene used once), with two-sided Mann-Whitney U tests and Benjamini-Hochberg correction across all twelve tests (**Supplementary Fig. 1g**).

#### Expression autocorrelation across cell divisions

Cells from clones with at least two cells were retained, and log-normalized expression was used throughout. Clones were binned by size into 2, 3-4, 5-8, 9-16, 17-32 and ≥33 cells, corresponding to a minimum of one, two, three, four, five and six cell divisions since the clone’s founding cell (a clone of n cells requires at least ⌈log_2_(n)⌉ divisions), and bins containing fewer than 15 clones were discarded. Within each bin, gene expression autocorrelation was quantified per gene as the one-way random-effects intraclass correlation coefficient^58^, ICC = (MSB − MSW) / (MSB + (n_0_ − 1) MSW), where MSB and MSW are the between-and within-clone mean squares and n_0_ = (N − Σ_k_ n_k_^2^ / N) / (K − 1) is the effective clone size for unequal clone sizes, with N cells, K clones and n_k_ cells in clone k. Genes without expression variance in a bin were left undefined. Because a clone of n cells has undergone approximately log_2_(n) divisions, each bin was placed at the mean log_2_ clone size of its cells. Consensus memory and non-memory genes were defined per group (called by at least two methods; see above), and the median and interquartile range of ICC across each gene set were plotted per bin. For Michaels and Weinreb, only the last time point is shown, because the earlier groups consist predominantly of two-cell clones.

#### Gene Ontology enrichment

Consensus memory genes (merged mode, called by at least two methods; see above) of each dataset were tested for over-representation of Gene Ontology^39,40^ terms by a one-sided Fisher’s exact test against that dataset’s own measured transcriptome, defined as all genes with non-zero mean expression; no expression floor was applied. P-values were Benjamini-Hochberg corrected^49^ across terms within each dataset, and terms with fewer than 10 or more than 500 genes in the background were excluded. Gene sets (Biological Process, Molecular Function and Cellular Component) were taken from MSigDB^59,60^ v2026.2: the C5 GO collection in human symbols for Michaels and Harmange, and the M5 GO collection in mouse symbols^61^ for Weinreb and Bandler, so that each dataset was tested in its own gene namespace. For the comparison across datasets, mouse genes were mapped to human symbols as described above (see Comparison across methods and datasets) and all four datasets were tested against the human collection, so that terms are comparable across species. Per-dataset and cross-dataset term counts therefore come from separate runs (all enriched terms are listed in **Supplementary Data S2**).

### Perturb-seq screen analysis

To test whether memory genes are more responsive to perturbation, we collected six published CRISPR perturbation screens with single-cell or bulk transcriptomic readout, spanning mouse and human, CRISPR knockout and CRISPRi knockdown modalities, and genome-scale and focused libraries, with cell types matched to those of the lineage-barcoded datasets (**Table S3**). All screens were processed through a single perturbation-response pipeline (see *Perturbation response quantification)*.

#### Genome-scale CRISPRi in primary human T cells

Zhu et al.^28^ generated a pooled CRISPRi screen with single-cell transcriptomic readout in primary human T cells from four healthy donors, each profiled in three activation states: unstimulated (Rest) and 8 hours and 48 hours after stimulation (Stim8hr, Stim48hr). The screen comprises 5,067,221 cells and 13,992 genes across the twelve donor x condition groups (GEO, GSE314342). Every group contains cells carrying non-targeting control guides (15,816-18,323 cells per group), and each donor received essentially the entire library, with 4,655 to 6,085 perturbations represented per donor against roughly 6,520 per condition. Perturbed cells were therefore pooled across all donors and activation conditions for the main analysis, and results computed separately within each donor and each activation condition are shown in **Supplementary Fig. 3c**. Retaining perturbations with at least 20 cells left 6,646 perturbations in the pooled analysis, and 6,514, 6,526 and 6,514 within the unstimulated, 8 h and 48 h conditions respectively. The 6,345 perturbations present in all three conditions were used for cross-condition comparisons on 1,601,952, 1,579,627 and 1,656,887 cells, respectively.

#### Arrayed CRISPR knockout of transcription factors in primary human CD4⁺ T cells

Weinstock et al.^30^ performed individual CRISPR knockouts of 84 transcription factors in primary human CD4⁺ T cells, each profiled by bulk RNA sequencing; the targets comprise transcription factors implicated in inborn errors of immunity and a set matched for constraint and expression level without known immune-disease association. We used the deduplicated count matrix (GEO, GSE271788), comprising 311 libraries from four donors: 252 knockout samples (each target in three donors) and 59 AAVS1-targeting control samples (8-20 per donor). The median library size was 2.1 million assigned reads. After mapping Ensembl identifiers to gene symbols with MyGene.info, the matrix contained 45,511 genes, of which 16,956 exceeded the control-expression floor (0.1 CPM; see Perturbation response quantification). Log fold changes (LFCs) of each knockout sample were computed against the control samples from the same donor, and these were averaged across donors per target. All 84 targets were retained.

#### Genome-scale in vivo CRISPR knockout across the mouse brain

Shi et al.^29^ generated an in vivo Perturb-seq screen of 1,947 disease-associated genes across the mouse brain. We obtained the released whole-brain CRISPR atlas from the authors’ Hugging Face repository (perturbai/wholebrain_crispr_atlas, revision a7c65dc, downloaded 16 July 2026). Cells assigned exactly one guide and passing the authors’ quality control were grouped by the authors’ predicted cell-type annotation into 27 populations, and log fold changes were computed within each population against its own non-targeting cells (214,698 in total), so that differences in cell-type composition between targets cannot contribute; safe-targeting guides were not used as controls. Dropping combinations of target and population with fewer than 20 cells, or with no non-targeting cells, left 3,421,749 cells and 19,070 genes in 44,160 target-by-population combinations, covering 1,946 targets.

#### CRISPRi in untreated melanoma cells

Xu et al.^31^ developed PerturbFate, a combinatorial-indexing single-cell platform that profiles chromatin accessibility, nascent and steady-state transcription, applied to CRISPR interference of more than 140 vemurafenib-resistance-associated genes in A375 melanoma cells. We used the RNA modality of the vehicle (DMSO) arm (GEO, GSE291147), matching the untreated state of the Harmange cells; the vemurafenib arm was not analyzed. Because few cells carried a single detected guide, cells were assigned to the most abundant guide if it accounted for at least half of the cell’s sgRNA UMIs and the cell had at least three sgRNA UMIs in total. The twelve non-targeting guides were combined into one control group. This gave 30,572 cells and 60,706 genes covering 142 targets, all with at least 20 cells (median 173 cells per target, range 24 to 748), together with 2,106 non-targeting control cells.

#### CRISPR knockout in patient-derived melanoma cells

Frangieh et al.^32^ developed Perturb-CITE-seq, which couples pooled CRISPR-Cas9 knockout to single-cell transcriptome and surface-protein readout, in patient-derived melanoma cells under three conditions: untreated, interferon-γ, and co-culture with autologous tumour-infiltrating lymphocytes. The screen targeted 248 genes associated with cancer-cell-intrinsic resistance to immune-checkpoint inhibitors with approximately 750 guides in approximately 218,000 cells. We used the untreated condition, matching the untreated state of the Harmange cells; the interferon-γ and co-culture conditions were not analyzed. Counts were taken from the harmonized scPerturb^62^ release of the original deposit (Single Cell Portal, SCP1064), downloaded from Zenodo (10.5281/zenodo.7041848, July 2026), in which guides are collapsed to their target gene, non-targeting cells are labelled as controls, and only cells with a single perturbation are retained; no further guide assignment was therefore applied. This gave 57,627 cells and 23,712 genes covering 248 targets, of which 235 had at least 20 cells (median 175 cells per target among those retained, range 20 to 408), and 15,361 control cells.

#### Focused CRISPRi in primary mouse haematopoietic progenitors

Bowness et al.^33^ (GEO, GSE320250) performed a pooled CRISPR interference screen with single-cell transcriptomic readout in primary mouse haematopoietic stem and progenitor cell (HSPC) expansion cultures, targeting 520 genes with three guides each, plus 120 non-targeting guides. Cells were profiled with Evercode Whole Transcriptome v3 (Parse Biosciences), with guides recovered by targeted amplification, in two independently initiated cultures: replicate 1 from five adult male mice (eight sublibraries) and replicate 2 from five further mice (three female, two male; three sublibraries). Guides were called at ≥2 guide UMIs, and cells assigned to a single target gene were retained, including cells with several guides against the same gene. This gave 95,848 cells (replicate 1, 79,287; replicate 2, 16,561) and 57,186 genes, covering 498 targets (median 131 cells per target, range 20 to 1,551) and 8,274 non-targeting control cells. LFCs were computed within each replicate, which is the biological batch here and is confounded with sex (see Perturbation response quantification). Sublibraries, aliquots of one barcoded cell pool split before sequencing, were not treated as a grouping variable.

**Table S3:** Perturb-seq screens paired with the lineage-barcoded datasets. The six perturbation screens used in **Fig. 2**, each paired with the lineage-barcoded dataset profiling the same cell type (**Table S1**). Library gives the number of targets in the published screen; perturbations analyzed are those retained after the 20-cell threshold. Cells are those with an assigned guide passing quality control, and are therefore fewer than the totals reported by the authors. Weinstock et al. is a bulk screen, so its count is sequencing libraries rather than cells, with control libraries in brackets. Where a screen has several treatment arms, only the untreated or vehicle arm was used, matching the untreated state of the lineage-barcoded cells. Fold changes were computed within each screen’s grouping variable (last column). Full screen metadata are in **Supplementary Data S4**.

| Dataset | Publication | Accession | Assay, modality | Cells perturbed | Library | Perturbations analyzed | Cells profiled | Paired lineage dataset |
|---|---|---|---|---|---|---|---|---|
| Zhu | Zhu et al. 2026, *Cell* 189(22):6988-7012.e17 | GSE314342 | probe-based Perturb-seq, CRISPRi | primary human CD4<sup>+</sup> T cells, 4 donors x 3 activation states | genome-scale | 6,646 | 5,067,221 (22 million in the paper) | Michaels |
| Weinstock | Weinstock et al. 2024, *Cell Genomics* 4(11):100671 | GSE271788 | arrayed CRISPR knockout, bulk RNA-seq | primary human CD4<sup>+</sup> T cells, 4 donors | 84 transcription factors | 84 | 311 libraries (252 knockout, 59 control) | Michaels |
| Shi | Shi et al. 2026, bioRxiv | perturbai/wholebrain_crispr_atlas | in vivo Perturb-seq, CRISPR knockout | mouse brain, 27 cell populations | 1,947 disease genes | 1,946 | 3,421,749(7.7 million in the paper) | Bandler |
| Xu | Xu et al. 2026, *Nature* 654(8117):261-271 | GSE291147 | PerturbFate, CRISPRi | A375 melanoma, DMSO arm | 142 resistance genes | 142 | 30,572 (>300,000 in the paper) | Harmange |
| Frangieh | Frangieh et al. 2021, *Nature Genetics* 53(3):332-341 | SCP1064 | Perturb-CITE-seq, CRISPR knockout | patient-derived melanoma, untreated arm | 248 genes, ~750 guides | 235 | 57,627 (~218,000 in the paper) | Harmange |
| Bowness | Bowness et al. 2026, bioRxiv | GSE320250 | Perturb-seq, CRISPRi | primary mouse HSPC cultures, 2 replicates | 520 genes | 498 | 93,146 (95,848 in the paper) | Weinreb |

#### Further Perturb-seq screens for the generalization analysis

To test whether the response result depends on the screens paired with our lineage datasets, we collected eight further published CRISPR screens with single-cell transcriptomics readout, covering interference and activation modalities, genome-scale and focused libraries, and eight human cell lines (K562, RPE1, HepG2, Jurkat, HCT116, HEK293T, TeloHAEC and the induced pluripotent line KOLF2.1J)^63–71^. Three of these screens (Swinderman et al.^71^, Wong et al.^68^, and Huang et al.^65^) are described in preprints that had not been reviewed at the time of writing. Screens whose design precludes a pooled analysis were divided into independent units: Wong et al.^68^ by treatment condition, as basal and cytokine-treated cells are different states, and Nourreddine et al.^67^ by library batch, as its library is split across batches so that fold changes must be computed within one. Publications reporting several screens contribute one unit per screen (three for Replogle et al.^63^, two each for Nadig et al.^64^ and Huang et al.^65^). This gives 15 analysis units comprising 68,659 perturbations and 13.3 million cells, of which 634,968 are unperturbed controls. Three screens contain paired perturbations in addition to single-gene perturbations^69–71^; within each, single and paired perturbations were analyzed as separate sets. Accessions, modality, library scale, perturbations analyzed, and cell numbers are given in **Table S4**.

**Table S4:** Perturb-seq screens used in generalization analysis. The 15 analysis units, from eight publications, on which Power-Seek was run in the control cells (**Fig. 2j, Supplementary Fig. 7**). An analysis unit is a set of cells processed independently end to end: pseudobulk profiles, fold changes, the Power-Seek run, and the expression-matched background are all computed within it. Publications reporting several screens contribute one unit per screen, and two were divided further because their design precludes pooling^67,68^. Perturbations are those retained after the 20-cell threshold, and are therefore fewer than the library sizes reported in the original publications. The paired column gives perturbations targeting two genes, which were analyzed as a separate set in the three screens that contain them. Cells are all cells of the unit passing quality control, with unperturbed control cells in brackets. Accessions identify the deposit from which the data were obtained. Full screen metadata, including download locations, file sizes, and the original library sizes, are given in **Supplementary Data S5**.

| Analysis unit | Publication | Accession /DOI | Assay, modality | Cell line | Perturbations | Paired | Cells (controls) |
|---|---|---|---|---|---|---|---|
| K562 essential | Replogle 2022 | Zenodo 7041849 | Perturb-seq, CRISPRi | K562 | 2,007 | – | 279,272 (10,311) |
| K562 genome-wide | Replogle 2022 | Zenodo 7041849 | Perturb-seq, CRISPRi | K562 | 9,728 | – | 1,805,466 (69,239) |
| RPE1 | Replogle 2022 | Zenodo 7041849 | Perturb-seq, CRISPRi | RPE1 | 2,064 | – | 213,461 (10,907) |
| HepG2 | Nadig 2025 | GSE264667 | Perturb-seq, CRISPRi | HepG2 | 2,070 | – | 129,925 (4,758) |
| Jurkat | Nadig 2025 | GSE264667 | Perturb-seq, CRISPRi | Jurkat | 2,197 | – | 232,054 (11,337) |
| HCT116 | Huang 2025 | Figshare 55021257 | FiCS, CRISPRi | HCT116 | 17,676 | – | 3,049,654 (148,576) |
| HEK293T | Huang 2025 | Figshare 55074802 | FiCS, CRISPRi | HEK293T | 17,938 | – | 4,141,479 (200,398) |
| Untreated | Wong 2026 | Zenodo 18792213 | Perturb-seq, CRISPRi | TeloHAEC | 1,560 | – | 384,544 (10,447) |
| IL-1β/TNFα treated | Wong 2026 | Zenodo 18792213 | Perturb-seq, CRISPRi | TeloHAEC | 1,574 | – | 389,560 (10,796) |
| Batch ALPHA | Nourreddine 2026 | Figshare 64650261 | Perturb-seq, CRISPRi | KOLF2.1J | 3,639 | – | 937,032 (78,387) |
| Batch BETA | Nourreddine 2026 | Figshare 64650261 | Perturb-seq, CRISPRi | KOLF2.1J | 3,857 | – | 930,743 (36,907) |
| Batch GAMMA | Nourreddine 2026 | Figshare 64650261 | Perturb-seq, CRISPRi | KOLF2.1J | 3,850 | – | 663,689 (25,149) |
| K562 | Norman 2019 | GSE133344 | Perturb-seq, CRISPRa | K562 | 105 | 131 | 102,922 (11,251) |
| K562 | Replogle 2020 | GSE146194 | direct-capture Perturb-seq, CRISPRi | K562 | 35 | 37 | 32,664 (4,296) |
| K562 | Swinderman 2026 | SRA SUB15796302 | ProPer-Seq, CRISPRa | K562 | 84 | 107 | 50,009 (2,209) |

### Perturbation response quantification

For the five single-cell screens, raw counts were summed over the cells of each perturbation within each group (defined below) to form pseudobulk profiles, which were normalized to counts per million (CPM); perturbations with fewer than 20 cells in a group were discarded in that group. The bulk screen from Weinstock et al.^30^ was normalized to CPM directly. LFCs were computed as LFC = log_2_((CPM_KO/KD_+1)/(CPM_ctrl_+1)), where CPM_ctrl_ is the pseudobulk of the control cells in the same group. Groups were the donor (Weinstock^30^), predicted cell type (Shi^29^), and biological replicate (Bowness^33^). Xu^31^ and Frangieh^32^ formed a single group each, as only their untreated arms were used, and Zhu^28^ was pooled across all four donors and all three activation conditions. Results computed within each donor and within each activation condition separately are shown in **Supplementary Fig. 3c**. For perturbations measured in several groups, per-group LFCs were averaged, weighted by the number of cells of that perturbation in each group (single-cell screens) or equally across donors (Weinstock, where each target was profiled in three of the four donors^30^). The LFC of each perturbation’s target gene was masked, as it reflects the perturbation itself rather than a transcriptional response.

Baseline expression of each gene was the mean expression over control cells, weighted across groups as for the LFCs. Genes were retained if their mean expression in control cells exceeded 0.1 UMI per cell (single-cell screens^28,29,31–33^) or 0.1 CPM (bulk-screen of Weinstock et al.^30^). Results for floors from no floor to 2 UMI per control cell, computed in Zhu, are shown in **Supplementary Fig. 3b**.

From the resulting perturbation x gene LFC matrix, we derived two readouts: the response of a gene, the mean |LFC| of that gene across all retained perturbations in which it was measured, excluding its own masked entry (**Fig. 2e-j**), and the impact of a perturbation, the mean |LFC| it causes across all retained genes (**Fig. 2b**).

### Response of memory genes

Each lineage dataset was paired with the screens from the same system: Michaels^23^ with Zhu^28^ and Weinstock (T cells)^30^, Harmange^17^ with Xu^31^ and Frangieh (melanoma)^32^, Weinreb^24^ with Bowness (HSPCs)^33^ and Bandler^25^ with Shi (brain)^29^.

#### Perturbation impact

Each perturbation was summarized by its impact, the mean |LFC| across all retained genes, excluding the perturbation’s own target. Perturbations of consensus memory genes (called by at least two methods within their genome-wide top N) were compared with all perturbations of genes called by no method. Memory genes were taken from the lineage dataset paired with each screen. Screens with fewer than ten memory-gene perturbations are shown without a memory-gene distribution or statistics, their individual knockouts plotted as points (Weinstock^30^, 1; Xu^31^, 5).

#### Background genes and expression matching

In every comparison, background genes are genes that none of the methods placed in their top N, with N the cutoff used for the memory set. Background genes were matched to the memory set on mean control expression: the expressed genes of the screen were divided into 50 quantile bins, and from each bin as many background genes were drawn as the memory set has in that bin, sampling with replacement where a bin contained too few. This was repeated 100 times. Within each draw, the background genes were ranked by response, and the 100 draws were averaged rank by rank, so that the control set has the same number of values as the memory set rather than 100 times as many. The resulting control values are therefore averages across draws rather than individual genes, and their spread is narrower than that of a single matched draw. Matching quality is reported as the standardized mean difference (SMD) in log10 expression between memory and control sets (all |SMD| < 0.10). Each pairing was analyzed in the expression space of its own screen, i.e., expressed genes, bins and response were all defined on the screen’s control cells. For every screen, each memory gene had non-memory genes available at comparable control expression (**Supplementary Fig. 3a**), so the matched comparison is not driven by a small number of repeatedly reused background genes.

#### Statistics

Memory and control sets were compared with two-sided Mann-Whitney U tests computed on each of the 100 matched draws. The reported p-value is the median across draws, with Benjamini-Hochberg correction^49^ across all tests in a panel. Effect sizes are reported as Cliff’s delta^72^ and as the percentage difference between medians.

#### Covariate matching

For consensus memory genes (at least two methods) from Michaels et al. in Zhu et al., controls were additionally matched on mean expression, detection rate, cell-state specificity and gene length, each in turn and all four jointly, by 1:1 nearest-neighbor matching^73^ on rank-standardized covariates: each memory gene was paired with one of its 10 nearest unused background genes, over 2,000 draws. Cell-state specificity is defined as log_2_ of the ratio between a gene’s highest and lowest mean control expression across the three activation states (Rest, Stim8hr and Stim48hr), with a pseudocount of 0.01.

#### Robustness to individual perturbations

Perturbations were ranked by their mean |LFC| across consensus memory genes and removed one at a time, strongest first, recomputing the mean |LFC| of the memory and control sets after each removal until 50 perturbations remained.

#### Pairings and consensus tiers

For each of the six pairings, memory sets were defined (i) per method, as its merged top 300 (**Fig. 2g**), and (ii) as consensus sets called by at least two, at least three, or all four methods (**Fig. 2h**). Estimates (percentage difference in median response) were summarized across pairings by their mean.

#### Sensitivity to cutoff and statistics

The per-method comparison was repeated at cutoffs of 100, 250, 500, 1,000, and 2,000 genes, for every method and every screen (**Supplementary Fig. 4b)**. Where a method ranked fewer genes than the cutoff it contributed all of them rather than being dropped; its memory set then stops growing with N while the background, the genes that no method placed in its top N, keeps shrinking, so the deeper cutoffs compare the same gene set against a progressively more restricted control pool. Response was replaced in turn by the median |LFC|, the standard deviation of LFC, the fraction of perturbations with |LFC| > 1 and the maximum |LFC|. As a negative control, the number of perturbations in which a gene was measured was analyzed in the same way.

#### Per-group memory sets

All perturbation analyses were repeated with memory sets defined within single experimental groups instead of merged across groups; for the impact comparison, groups with fewer than ten memory-gene perturbations were excluded.

#### Comparison with bulk MemorySeq

Memory genes of the Harmange dataset were compared with the 227 MemorySeq genes reported in bulk WM989 cells (Shaffer et al. 2020). Overlap was tested against 1,000 random gene sets of equal size drawn within 20 expression-rank bins of the genes measured in both assays. The response of both sets was compared in Xu and Frangieh, each against its own expression-matched control.

### Lineage-free memory genes

Power-Seek, GEMLI in lineage-free mode, and isCORAL^41^ were run on the unperturbed control cells of the five single-cell screens^28,29,31–33^, separately within each group of a screen. Weinstock et al.^30^ is a bulk screen and has no cell-level data. All three received the same exported control-cell matrices (log-normalized counts, genes above 0.1 UMI per control cell, the floor of the response analysis). Power-Seek and GEMLI were run with the settings used for the lineage-barcode datasets.

isCORAL, described in a preprint that had not been reviewed at the time of writing (package v1.0.0-beta, commit f9fa160, installed from the authors’ repository and used unmodified), applies CORAL’s w^2^ statistic to pseudoclones inferred from expression rather than to barcoded clones. It was run on the same matrix as the other callers, with the bin size set to N/50 as the package recommends and at most 4,000 cells per group for tractability. It returns a score rather than a call, so genes were ranked by w^2^, and those the package leaves unscored were dropped.

Per-group lists were merged by best rank, as for the lineage datasets. The top 300 genes of each method, and the genes called by all three on the control cells, were tested for response in the same screen against an expression-matched control, and each control-cell set is shown next to the same method run with barcodes on the cell-type-matched lineage datasets. isCORAL’s barcode counterpart is CORAL. These sets were not used in any cross-method analysis.

### Generalization across Perturb-seq screens

For each of the 15 analysis units, perturbation profiles were built exactly as above: raw counts were summed over the cells of each perturbation to form pseudobulk profiles, perturbations with fewer than 20 cells were discarded, profiles were normalized to counts per million, and LFCs were computed against the unperturbed control cells of the same screen as LFC = log_2_((CPM_pert + 1)/(CPM_ctrl + 1)). In the three screens with paired perturbations, guide variants of the same target were merged, and labels were canonicalized to the sorted set of targets, and every target of a label was masked in its own row. Genes were retained if their mean expression in the control cells exceeded 0.1 UMI per cell, the same criterion as for the main screens. Memory genes were the top 300 of a Power-Seek run on that screen’s own unperturbed control cells, with no lineage or clone information, restricted to the retained genes. The background was an equal-sized, expression-matched set drawn from the genes Power-Seek did not call, using 50 quantile bins and 100 draws as above. The response was compared with two-sided Mann-Whitney U tests, Benjamini-Hochberg corrected across all 18 comparisons: the 15 analysis units and the paired-perturbation sets of the three screens that contain them^69–71^. Effect sizes are reported as Cliff’s delta and as the percentage difference between medians (**Fig. 2j**; per-screen values in **Supplementary Fig. 7**).

### Statistics and reproducibility

All statistical tests were two-sided unless stated otherwise. Comparisons of gene sets used two-sided Mann-Whitney U tests. Effect sizes are reported as Cliff’s delta, the probability that a memory gene exceeds a control gene minus the reverse (range-1 to 1), and as the percentage difference between medians. P-values were Benjamini-Hochberg corrected within each figure panel, or within each row for the grid panels, except where stated. Resampling procedures (random pairing, permutation, expression matching, and bootstrap) used fixed random seeds, and the number of iterations is given with each analysis. No statistical method was used to predetermine sample size. All analyses used published datasets, and no data were excluded except as described.

### Software

All analyses other than the memory-gene callers were carried out in Python 3.10.20 with numpy 2.2.6, scipy 1.15.2, pandas 2.3.3, anndata 0.11.4, scanpy 1.11.5, statsmodels 0.14.6, and scikit-learn 1.7.2. Figures were produced with matplotlib 3.10.9 and seaborn 0.13.2. Ortholog and synonym resolution additionally used mygene 3.2.2. CORAL and GEMLI were run under R 4.4.3. CORAL used Seurat 5.5.1, Matrix 1.7.6, and ComplexHeatmap 2.22.0. GEMLI required only base R and stats when clone labels were supplied, and dendextend and igraph in its lineage-free mode. Power-Seek was run in Python from the authors’ notebooks. The commit of each published caller is given with its description above.

