## Supplementary Tables and Information for "Memory genes respond strongly to genetic perturbations and have CpG-depleted promoters": SI_Mathematical_response_model_BKS.docx

### **Supplementary Methods —**

### **Model for separable Rank-1 RMS responses**

Consider a system of $n$ genes with expression state $\mathbf{x}\in\mathbb{R}^{n}$ and dynamics $\dot{\mathbf{x}}=\mathbf{F}\left( \mathbf{x} \right)+\mathbf{u}^{\left( i \right)}$. Here $i\in\{1,\ldots,P\}$ indexes the $P$ distinct perturbations, $\mathbf{u}^{\left( i \right)}\in\mathbb{R}^{n}$ is the effective input induced by perturbation $i$, and $j\in\{1,\ldots,n\}$ indexes target genes. At a stable reference state $\mathbf{x}_{0}$, let $J_{jk}=\left. \partial F_{j}/\partial x_{k} \right|_{\mathbf{x}_{0}}$ denote the full regulatory Jacobian. For weak, constant inputs and invertible $J$, the steady-state response is

$\delta\mathbf{x}^{\left( i \right)}\equiv\mathbf{x}^{\left( i \right)}-\mathbf{x}_{0}\simeq-J^{-1}\mathbf{u}^{\left( i \right)}.$	(S1)

No block-diagonal or rank-one assumption is imposed on $J$. This derivation only assumes perturbations share an approximately common reference state and Jacobian.

**Measured CPM log-fold changes.** Define the total RNA count $T\left( \mathbf{x} \right)=\sum_{k=1}^{n} x_{k}$, CPM expression $c_{j}\left( \mathbf{x} \right)=Kx_{j}/T\left( \mathbf{x} \right)$ with $K={10}^{6}$, and transformed observable $y_{j}\left( \mathbf{x} \right)=log\left[ 1+c_{j}\left( \mathbf{x} \right) \right]$. The measured response to perturbation $i$ is $\mathcal{l}_{ij}=y_{j}\left( \mathbf{x}^{\left( i \right)} \right)-y_{j}\left( \mathbf{x}_{0} \right)=log\{\left[ 1+c_{j}\left( \mathbf{x}^{\left( i \right)} \right) \right]/\left[ 1+c_{j}\left( \mathbf{x}_{0} \right) \right]\}$. Let $T_{0}=T\left( \mathbf{x}_{0} \right)$ and $c_{0j}=c_{j}\left( \mathbf{x}_{0} \right)$. The measurement Jacobian $B\in\mathbb{R}^{n\times n}$ has entries

$B_{jk}=\left. \frac{\partial y_{j}}{\partial x_{k}} \right|_{\mathbf{x}_{0}}=\frac{K}{T_{0}\left( 1+c_{0j} \right)}\left( \delta_{jk}-\frac{x_{0j}}{T_{0}} \right).$	(S2)

Equivalently, $B=D\left( K/T_{0} \right)\left[ I-\mathbf{x}_{0}\mathbf{1}^{T}/T_{0} \right]$, where $D=diag\left[ \left( 1+c_{01} \right)^{-1},\ldots,\left( 1+c_{0n} \right)^{-1} \right]$. The effective observed-response Jacobian is $A\equiv-BJ^{-1}$, so that

$\mathcal{l}^{\left( i \right)}\simeq B\delta\mathbf{x}^{\left( i \right)}=A\mathbf{u}^{\left( i \right)}, \mathcal{l}_{ij}\simeq\sum_{k=1}^{n} A_{jk}u_{k}^{\left( i \right)}.$	(S3)

For base-2 LFCs, $B$ and $A$ are divided by $\ln2$. Thus $A$ includes intrinsic dynamical susceptibility, CPM compositional normalization, and the pseudocount transformation.

**RMS response and source–target separation.** Let $p\in\{S,F\}$ index the source class of perturbed genes and $q\in\{S,F\}$ the target class of measured genes. Write $\mathcal{I}_{p}\subseteq\{1,\ldots,P\}$ for perturbations in source class $p$, with $P_{p}=\left| \mathcal{I}_{p} \right|$, and $\mathcal{J}_{q}\subseteq\{1,\ldots,n\}$ for target genes in class $q$, with $N_{q}=\left| \mathcal{J}_{q} \right|$. Define the target-selection matrix $\Pi_{q}\in\{0,1{\}}^{N_{q}\times n}$, whose rows select the coordinates $j\in\mathcal{J}_{q}$. (We reserve $P$ for the number of perturbations.) For complete measurements, the pooled source–target RMS is

$L_{pq}^{2}\equiv\frac{1}{P_{p}N_{q}}\sum_{i\in\mathcal{I}_{p}} \sum_{j\in\mathcal{J}_{q}} \mathcal{l}_{ij}^{2}=\frac{1}{P_{p}N_{q}}\sum_{i\in\mathcal{I}_{p}} \parallel\Pi_{q}\mathcal{l}^{\left( i \right)}\parallel^{2}.$	(S4)

Define the uncentered second-moment matrix of inputs from $p$ as $Q_{p}\equiv P_{p}^{-1}\sum_{i\in\mathcal{I}_{p}} \mathbf{u}^{\left( i \right)}\mathbf{u}^{\left( i \right)T}$; it is not the mean-subtracted covariance unless the mean input vanishes. Substitution of the linear response gives

$L_{pq}^{2}\simeq\frac{1}{N_{q}}\mathrm{Tr}\left( \Pi_{q}AQ_{p}A^{T}\Pi_{q}^{T} \right).$	(S5)

A sufficient condition for separability is that source classes excite the same normalized input second-moment structure, differing only in overall RMS amplitude:

$Q_{p}\simeq G_{p}^{2}Q, p\in\{S,F\},$	(S6)

for one positive-semidefinite matrix $Q$ independent of $p$. It follows that

$L_{pq}\simeq G_{p}H_{q}, H_{q}^{2}\equiv\frac{1}{N_{q}}\mathrm{Tr}\left( \Pi_{q}AQA^{T}\Pi_{q}^{T} \right).$	(S7)

Neither $Q$ nor $A$ need be isotropic or rank one. The condition on $Q_{p}$ is sufficient, but not necessary.

**Predictions and distinction from gene-level rank one.** For two source and two target classes, the RMS matrix $\left[ L_{pq} \right]$ is approximately rank one when

$C\equiv\frac{L_{SS}L_{FF}}{L_{SF}L_{FS}}\simeq1, \frac{G_{S}}{G_{F}}\simeq\sqrt{\frac{L_{SS}L_{SF}}{L_{FS}L_{FF}}}, \frac{H_{S}}{H_{F}}\simeq\sqrt{\frac{L_{SS}L_{FS}}{L_{SF}L_{FF}}}.$	(S8)

Separation of $Q_{p}$ alone does not imply that the raw LFCs are rank one. Similarly, a nearly rank-one matrix of group RMS amplitudes does not imply a rank-one signed, gene-resolved LFC matrix. Finally, the approximation assumes a valid local linearization of both the dynamics and CPM-log measurement: the inferred $H_{q}$ is susceptibility of the normalized observable, not that including absolute RNA abundance. Large state shifts may break linear response as well as separability.

Figure B | Source potency, target susceptibility, and RMS factorization

We divided the perturbations and measured genes in each dataset into memory (S) and nonmemory (F) classes. Memory sets were obtained from dataset-specific ranked gene lists (up to 300 genes per list), and perturbation and target identities were matched to the LFC matrix. For source perturbation class p and target gene class q, we calculated the pooled RMS of the signed log-fold-change responses over all finite entries in the corresponding source–target block. The four RMS values L_SS, L_SF, L_FS, and L_FF define a 2 × 2 matrix of source–target amplitudes. Under the separable the factorization constant C equals one, while ratios of the source potencies and target susceptibilities can be estimated algebraically.

To control for unequal source and target group sizes, we used 300 bootstrap draws per dataset. In each draw, the memory and nonmemory source groups were sampled with replacement to the smaller source-group size; memory targets were sampled with replacement and nonmemory targets without replacement to the smaller target-group size. We computed the four block RMS values and the derived C, G_S/G_F, and H_S/H_F ratios for each draw, excluding draws with missing or nonpositive block RMS values. Dataset-level estimates are bootstrap medians, and error bars show the 2.5th–97.5th percentiles of the bootstrap draws.

Twenty-three of 24 total datasets had sufficiently represented memory and nonmemory source and target classes. In panel B, translucent violins show the distribution of dataset-level estimates, colored points show individual datasets, light vertical lines show bootstrap percentile intervals, and black diamonds with error bars indicate the across-dataset mean ± SEM. Horizontal dashed lines mark unity. The panel is ordered as the factorization constant C (gray), source-potency ratio G_S/G_F (blue), and target-susceptibility ratio H_S/H_F (red). Tests of dataset-level values against unity use two-sided one-sample t-tests.

**Figure C | Dominance of the leading response mode**

We quantified the extent to which perturbation responses can be represented by a single collective mode, contrasting signed gene-level log-fold changes (LFCs) in memory and matched nonmemory genes with the unsigned root-mean-square (RMS) responses of target-gene populations. For each response matrix A (perturbations × measured features), the singular-value decomposition is A = UΣVᵀ. The fraction of squared response amplitude captured by its leading mode is

f₁ = σ₁² / Σₖ σₖ² . (S13)

Here σ₁ is the largest singular value; f₁ = 1 indicates an exactly rank-one matrix, whereas smaller values imply appreciable higher-dimensional structure.

For the signed-LFC comparison, each of the 24 datasets supplied a perturbation-by-target-gene matrix. Self-perturbation entries, nonfinite measurements and |LFC| > 100 were excluded; for genes we required ≥70% finite coverage and non-negligible variation. Missing entries were imputed using the target gene’s median response. Up to 300 memory genes were retained per dataset (minimum 15), alongside 20 independently selected, equally sized nonmemory control sets matched without replacement by proximity in log RNA half-life. Each target-gene column was centered across perturbations, so that f₁ measures the leading fraction of between-perturbation LFC variation. The control statistic is the median f₁ across its 20 matched draws. Singular values were obtained numerically using a leading-singular-value algorithm, with full SVD as a fallback.

For total RMS, we assembled a positive, uncentered perturbation-by-population response matrix with six columns (Memory and Q1–Q5). The observed RMS of each target population was used, with the Q1 anchor reconstructed from the held-out prediction and the relevant population scaling factor.

The leading RMS fraction was then calculated from the squared singular values of this uncentered matrix. Because its entries are strictly positive and are not mean-centered, a large first-mode fraction reflects strong multiplicative/common-amplitude structure.

Panel C includes all 24 datasets, and each thin gray line joins the matched-nonmemory LFC, memory LFC, and total RMS first-mode fractions within a single dataset. Gray points denote individual datasets (with small horizontal jitter), while black diamonds and error bars show the across-dataset mean ± standard error of the mean (SEM; sample SD/√24). The plot shows relatively modest first-mode fractions for centered LFCs (typically below 0.35), with memory genes generally displaying somewhat greater leading-mode coherence than half-life-matched controls. By contrast, the uncentered total-RMS matrices have first-mode fractions near unity. This suggests a highly coherent population-level RMS amplitude across perturbations even when the underlying signed, gene-resolved changes are substantially higher-dimensional.

### **Figure D | Predicting gene responses across perturbations**

We analyzed 24 Perturb-seq datasets. For each dataset, we divided the measured genes into six groups: one group of memory genes (165–492 genes per dataset) and five approximately equal-sized groups of nonmemory genes (Q1–Q5). The nonmemory genes were ordered by RNA half-life, when available, with baseline expression or response variability used as a fallback. The size of each nonmemory group varied among datasets; Q1–Q5 were not separately matched control groups.

For each perturbation $i$, we measured the overall response of gene group $q$ using the root-mean-square (RMS) log-fold change:

$$R_{iq}=\left[ \frac{1}{N_{iq}}\sum_{j\in q_{i}} L_{ij}^{2} \right]^{1/2}. \text{(S9)}$$

Here $L_{ij}$ is the log-fold change of gene $j$ following perturbation $i$, and $N_{iq}$ is the number of genes with available measurements in that group. Under the rank-one model, the ratio of responses between any two gene groups should be approximately the same for different perturbations:

$$\frac{R_{iq}}{R_{ia}}\simeq\frac{R_{jq}}{R_{ja}}, i\neq j. \text{(S10)}$$

We tested this prediction using four-fold cross-validation within each dataset. Using three folds of perturbations, we calculated the geometric-mean ratio between the response of group $q$ and the reference group Q1:

$$\hat{r}_{q}^{\mathrm{train}}=exp\left[ \frac{1}{N_{\mathrm{train}}}\sum_{j\in train} \log\left( \frac{R_{jq}}{R_{j,Q1}} \right) \right]. \text{(S11)}$$

For each held-out perturbation $i$, we multiplied this ratio by its measured Q1 response to predict the response in group $q$:

$$\hat{R}_{iq}=R_{i,Q1} \hat{r}_{q}^{\mathrm{train}}. \text{(S12)}$$

We repeated the procedure until each perturbation had been held out once. Each point in Panel D compares the measured response $R_{iq}$ (horizontal axis) with the predicted response $\hat{R}_{iq}$ (vertical axis) for one held-out perturbation and one gene group. Each eligible perturbation contributed five points (memory genes and Q2–Q5); Q1 itself was excluded because it served as the measured reference. Agreement with the diagonal indicates that relative responses between gene groups remain approximately constant across perturbations, as expected under the rank-one model.

Figure E | Hierarchically clustered perturbation-response covariance

To investigate whether distinct genetic perturbations elicit shared transcriptional responses, we analyzed the covariance between their gene-expression response profiles. Two perturbations can therefore generate similar responses if they excite overlapping regulatory modes, even when they act on different genes. Shared slow or strongly amplified modes can produce extended blocks of positively or negatively covarying perturbation profiles.

We used the bowness_velten Perturb-seq dataset and constructed a signed log-fold-change (LFC) matrix X, with rows representing individual perturbations i and columns representing measured target genes g. The analysis retained genes observed in at least 90% of perturbations and perturbations with at least 90% usable targets, removed near-constant genes (standard deviation ≤ 10⁻⁸), and replaced remaining missing values with the median LFC for each gene. If more than 1,500 perturbations survived filtering, the 1,500 perturbations with the largest uncentered response RMS were selected. The analyzed perturbation set thus may be enriched for relatively strong responses.

To identify groups with shared response profiles, we separately normalized the covariance to Pearson correlation and defined the pairwise clustering distance

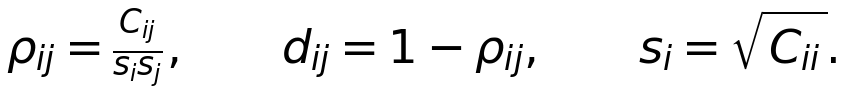
 (S15)

We applied average-linkage agglomerative hierarchical clustering to this distance matrix and reordered the rows and columns of C using the resulting dendrogram. We chose eight flat clusters to delineate block boundaries, which determines the black separators but not the hierarchical ordering. The displayed heatmap uses a red–white–blue scale centered at zero, with symmetric limits set by the 99th percentile of absolute covariance values to limit the effect of outliers.

In panel E, extensive off-diagonal structure suggests that responses are organized into recurring regulatory programs rather than varying independently across perturbations. Such blocks are consistent with low-dimensional collective modes in the perturbation-response operator.

Figure F | Spectral scaling of memory-gene responses

In the 24 Perturb-seq datasets we assembled a perturbation-response matrix X, where X_ig is the signed log-fold change (LFC) of target gene g under perturbation i. Gene profiles were centered and standardized across perturbations. For each subset of m target genes, we computed the gene–gene correlation matrix and the ratio of its leading eigenvalue to the median remaining (“bulk”) eigenvalue:

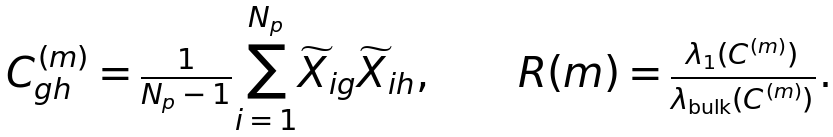
 (S13)

For independent standardized responses, R(m) is approximately unity (up to finite-sample fluctuations). A coherent rank-one contribution C ≈ I + α11ᵀ instead predicts a leading eigenvalue ≈ 1 + αm and bulk eigenvalues ≈ 1, so that R(m) ≈ 1 + αm. Thus, the increase in R with gene-set size measures the relative strength of shared response structure.

We evaluated R(m) on increasing gene-set sizes (starting at m = 12, in increments of 20, up to approximately 150 genes where coverage allowed) using dataset-specific ranked memory-gene lists. As a baseline, we generated 49 independently sampled nonmemory control sets per dataset, matching memory genes one-to-one without replacement by RNA half-life where available. The blue symbols show the median matched-control ratio and the blue bands its 2.5th–97.5th percentile range. Straight lines show ordinary least-squares fits R(m) = b + sm.

In ProPerSeq K562 CRISPRa, the memory-gene slope was sₘ = 2.135 versus sꜰ = 0.622 for matched nonmemory genes, a 3.43-fold enrichment. The dominant spectral component becomes increasingly separated from the remaining eigenvalues as additional memory genes are included. These findings support a low-dimensional collective response structure in memory genes for this dataset.

Figure G | Memory vs non-memory spectral scaling slopes

Across 24 perturbation datasets, we compared the spectral scaling of memory genes with matched nonmemory controls. For each dataset, we fitted a linear relationship between the leading-to-bulk eigenvalue ratio and the number of target genes, as illustrated in Fig. F:

*λ₁(m) / λ_bulk(m) = b + sm* (S14)

Here, s quantifies the rate at which the dominant collective response mode becomes spectrally separated as additional genes are included. Larger slopes indicate stronger collective organization of perturbation responses.

For each dataset, we calculated one memory-gene slope and the median slope across independently sampled matched-nonmemory control sets. This produced 24 paired comparisons. Gray points represent individual datasets, with connecting lines preserving the pairing between memory and nonmemory measurements. Black diamonds show the cross-dataset mean, with error bars representing the standard error of the mean (SEM).

Memory genes exhibited significantly larger spectral scaling slopes than matched nonmemory genes (two-sided paired t-test, n = 24, p = 0.021). The test was performed across dataset-level pairs, rather than treating control resamples as independent observations. The difference indicates that the dominant collective mode becomes increasingly pronounced with memory-gene set size, beyond the corresponding scaling observed in nonmemory controls.

These results support the hypothesis that memory genes are preferentially organized into coherent, low-dimensional regulatory response modes. Such organization is consistent with enhanced susceptibility to perturbations along collective directions of the gene regulatory network, although the spectral analysis alone does not establish the dynamical timescales of these modes.
