## Supplementary Figures for "Memory genes respond strongly to genetic perturbations and have CpG-depleted promoters"

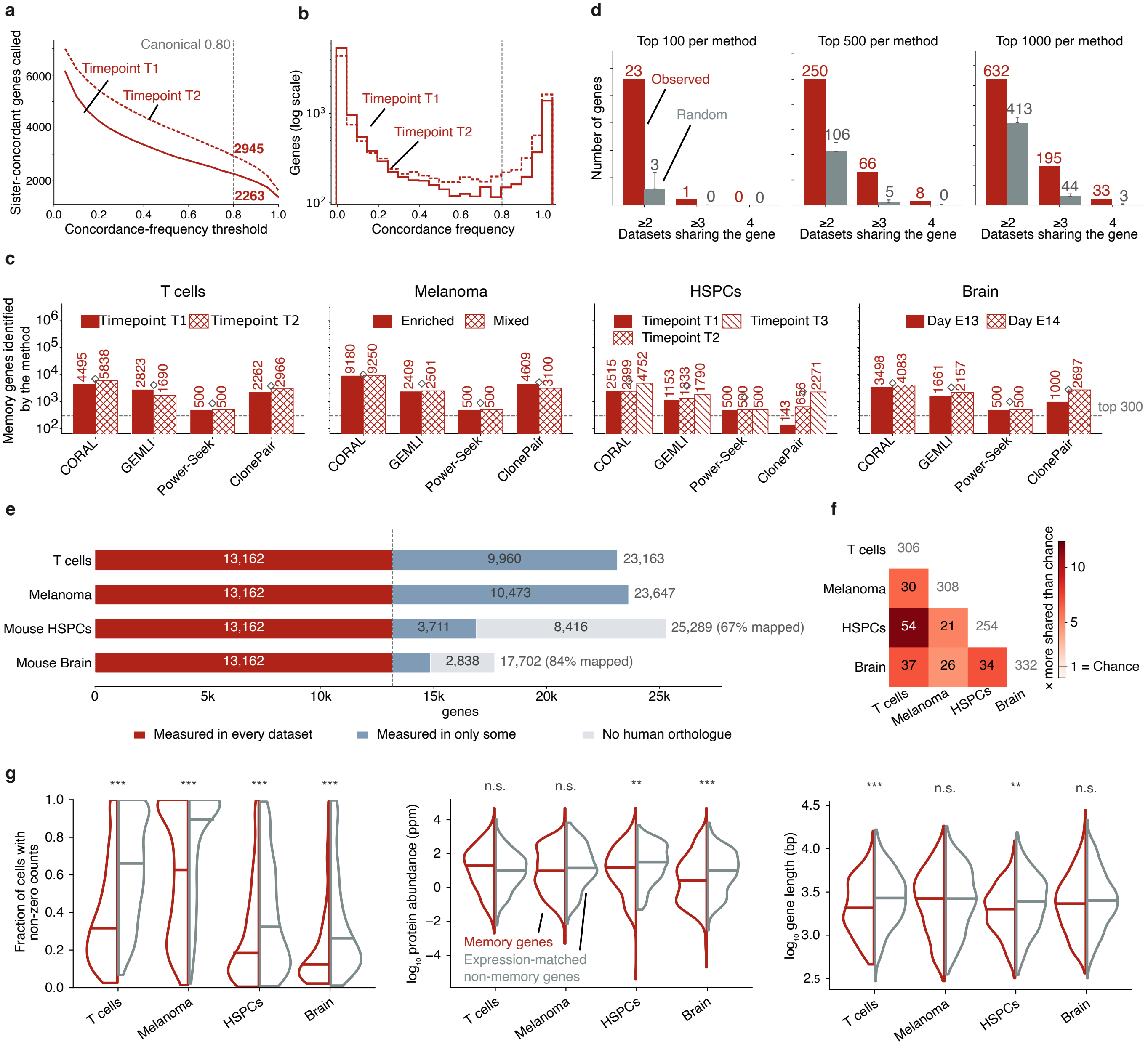
a
Canonical 0.80
Timepoint T1
Timepoint T2
2945
2263
Sister-concordant genes called
Concordance-frequency threshold
b
Timepoint T1
Timepoint T2
Genes (log scale)
Concordance frequency
d
Top 100 per method
Top 500 per method
Top 1000 per method
Observed
Random
Number of genes
Datasets sharing the gene
c
T cells
Melanoma
HSPCs
Brain
Timepoint T1
Timepoint T2
Enriched
Mixed
Timepoint T3
Day E13
Day E14
Memory genes identified by the method
top 300
CORAL
GEMLI
Power-Seek
ClonePair
e
T cells
Melanoma
Mouse HSPCs
Mouse Brain
13,162
9,960
23,163
10,473
23,647
3,711
8,416
25,289 (67% mapped)
2,838
17,702 (84% mapped)
genes
Measured in every dataset
Measured in only some
No human orthologue
f
× more shared than chance
1 = Chance
g
Fraction of cells with non-zero counts
log10 protein abundance (ppm)
log10 gene length (bp)
Memory genes
Expression-matched non-memory genes
n.s.

**Supplementary Fig. 1: The memory gene definition is robust with respect to its threshold, its depth and the shared gene universe.**
**a** Sensitivity of the ClonePair call to its concordance threshold, in T cells (Michaels[22], progenitor T cells). Genes are called from all within-clone pairs of clones of two to eight cells, the window used throughout. Number of genes called against the concordance-frequency threshold for timepoint 1 (solid) and timepoint 2 (dashed); line, mean over three independent reshuffles; band, their range; grey dashed vertical line, the canonical threshold of 0.80 used by the pipeline, with the count at that threshold given beside each curve.
**b** Distribution of the continuous concordance frequency underlying the call in **a**, for one representative reshuffle, for timepoint 1 (solid) and timepoint 2 (dashed). Gene counts on a logarithmic axis; grey dashed vertical line, the canonical threshold of 0.80.
**c** Ranking depth available per method, as the number of genes each method ranks per dataset and experimental group. Squares, the pooled value across groups; dashed line, the top 300 cutoff used in **Fig. 1**. Power-Seek[21] has a fixed ceiling of 500 genes per group.
**d** Cross-dataset sharing of memory genes at three further depths, **Fig. 1d** being the same analysis at the top 300. Per dataset, memory genes are those called by at least two of the four methods within each method's top N, with groups pooled into one ranking by best within-group rank. Mouse genes were mapped to human symbols, and all sets restricted to the 13,162 orthologs measured in every dataset. Red, observed number of genes shared by at least two, at least three, or all four datasets; grey, an expression-matched null in which each dataset's set is redrawn within its own expression-rank bins (20 bins, 1,000 permutations); bar height, permutation mean; whisker, 97.5th percentile. Each panel has its own y-axis, so bar heights compare within a panel only.
**e** Composition of each dataset's measured genes. Bars split the genes measured in a dataset into those measured in every dataset after ortholog mapping (13,162, the shared universe used in **Fig. 1d**), those carrying a human symbol but measured in only some datasets, and those without a human orthologue. Mouse datasets were mapped with NCBI's[33] curated ortholog table; the percentage of measured genes carrying a human symbol is given for each.
**f** Pairwise overlap of memory genes between datasets. Memory genes are those called by at least two of the four methods within each method's top 300, mapped to human symbols. Each pair is evaluated in its own universe of genes measured in both datasets (13,856 to 16,329 genes). Numbers off the diagonal, the count of shared memory genes; diagonal, each dataset's set size in that universe; colour, the ratio of the observed overlap to the overlap expected by chance under a hypergeometric model in the same universe, with 1 marking chance.
**g** Properties of memory genes other than expression, against an expression-matched background. Red, genes called by at least two of the four methods within each method's top 300; grey, genes no method called, matched one-to-one to the memory genes by nearest neighbour on log mean expression (k=10, each background gene used once). Panels, single-cell detection rate, protein abundance (PaxDb[34] integrated human, ppm), and exonic gene length (bp). Horizontal lines, medians; two-sided Mann-Whitney U test with Benjamini-Hochberg correction across all 12 tests (n.s., $q > 0.05$; ** $q < 0.01$; *** $q < 0.001$).

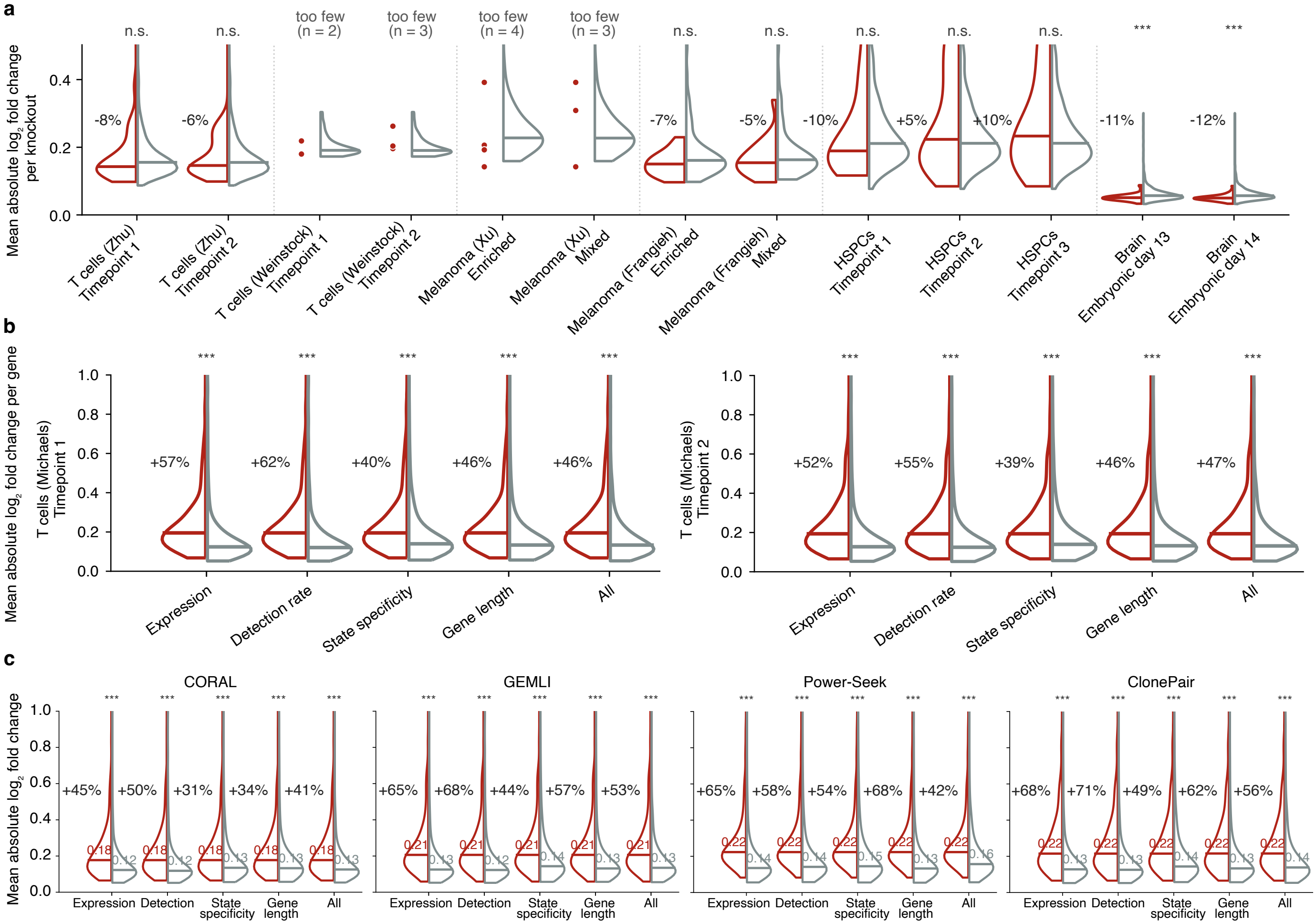

a
Mean absolute log2 fold change per knockout
n.s.
too few (n = 2)
too few (n = 3)
too few (n = 4)
***
-8%
-6%
-7%
-5%
-10%
+5%
+10%
-11%
-12%
T cells (Zhu) Timepoint 1
T cells (Zhu) Timepoint 2
T cells (Weinstock) Timepoint 1
T cells (Weinstock) Timepoint 2
Melanoma (Xu) Enriched
Melanoma (Xu) Mixed
Melanoma (Frangieh) Enriched
Melanoma (Frangieh) Mixed
HSPCs Timepoint 1
HSPCs Timepoint 2
HSPCs Timepoint 3
Brain Embryonic day 13
Brain Embryonic day 14
b
Mean absolute log2 fold change per gene
T cells (Michaels) Timepoint 1
T cells (Michaels) Timepoint 2
+57% +62% +40% +46% +46%
+52% +55% +39% +46% +47%
Expression
Detection rate
State specificity
Gene length
All
c
Mean absolute log2 fold change
CORAL
GEMLI
Power-Seek
ClonePair
+45% +50% +31% +34% +41%
+65% +68% +44% +57% +53%
+65% +58% +54% +68% +42%
+68% +71% +49% +62% +56%
Expression
Detection
State specificity
Gene length
All

**Supplementary Fig. 2: Memory genes respond more strongly under every confounder-matching scheme and in every experimental group, but perturbing them is no more consequential.**

**a** Transcriptome-wide impact of perturbations of memory genes, as in **Fig. 2c**, but with memory genes defined within each experimental group: a gene counts as a memory gene only if at least two of the four methods rank it in their top 300 for that group, with no merging across groups. Red, perturbations of those genes; grey, all perturbations of genes that no method called in that group. Each column is one combination of screen and lineage group, dotted lines separate screens. Groups with fewer than ten memory gene perturbations are shown without a memory gene distribution or statistics; their individual perturbations are plotted as points, with the counts above T cells (Weinstock[30]) and Melanoma (Xu[31]). Horizontal lines, medians; percentages, the difference between medians; asterisks, two-sided Mann-Whitney U test with Benjamini-Hochberg correction across the tested comparisons in each row (n.s., $q > 0.05$; **, $q < 0.01$).

**b** Response under confound matching, computed separately within each timepoint of the T cell dataset (Michaels[22]), otherwise as in **Fig. 2f**. Each row is one timepoint; each column a control set matched on mean expression, single-cell detection rate, cell-state specificity, gene length, or all jointly. Red, memory genes called by at least two methods within that timepoint's top 300; grey, matched controls; percentage, the increase in median over the control; asterisks, two-sided Mann-Whitney U test with Benjamini-Hochberg correction across the five comparisons of each row (*** $q < 0.001$).

**c** Each panel is one memory caller. Its top 300 memory genes, with timepoints 1 and 2 pooled into one ranking by best within-group rank, are compared with control sets matched on mean expression, single-cell detection rate, cell-state specificity, gene length, or all jointly, in the T cell screen (Zhu[27]). Controls are drawn from the genes which no method placed in its top 300. Expression is matched by quantile binning (50 bins, averaged over 100 draws); the other schemes by nearest-neighbour matching (k=10, 2,000 draws). All comparisons are restricted to genes above 0.1 UMI in that screen. Red, memory genes; grey, matched controls; number inside each half, medians; percentages, the increase in median over the matched control; asterisks, two-sided Mann-Whitney U test with Benjamini-Hochberg correction across the five schemes of each panel (*** $q < 0.001$). Distributions are truncated at 1.0 for display.

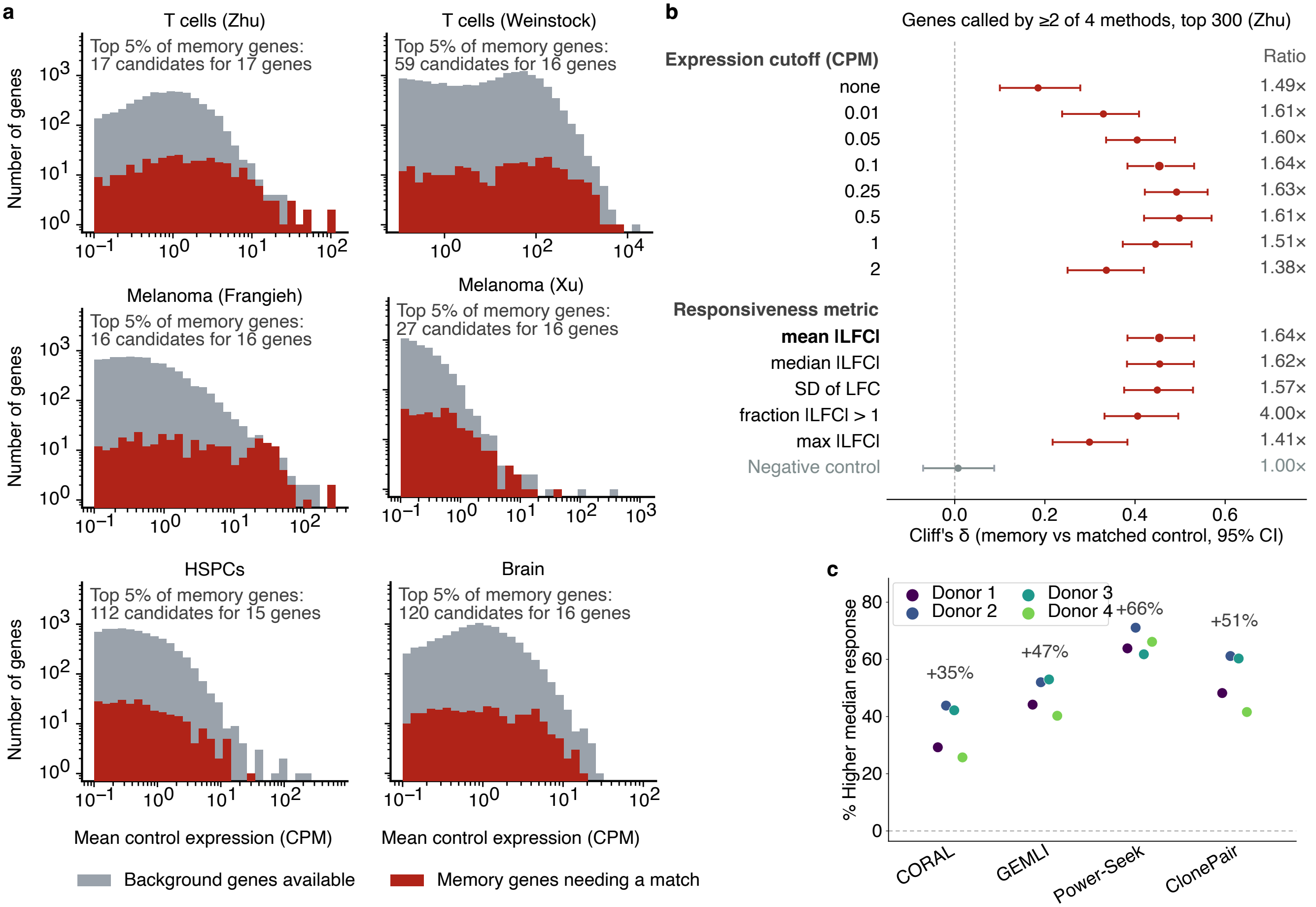
a
T cells (Zhu)
Top 5% of memory genes:
17 candidates for 17 genes
T cells (Weinstock)
Top 5% of memory genes:
59 candidates for 16 genes
Melanoma (Frangieh)
Top 5% of memory genes:
16 candidates for 16 genes
Melanoma (Xu)
Top 5% of memory genes:
27 candidates for 16 genes
HSPCs
Top 5% of memory genes:
112 candidates for 15 genes
Brain
Top 5% of memory genes:
120 candidates for 16 genes
Number of genes
Mean control expression (CPM)
Background genes available
Memory genes needing a match
b
Genes called by ≥2 of 4 methods, top 300 (Zhu)
Expression cutoff (CPM)
Ratio
none 1.49×
0.01 1.61×
0.05 1.60×
0.1 1.64×
0.25 1.63×
0.5 1.61×
1 1.51×
2 1.38×
Responsiveness metric
mean |LFC| 1.64×
median |LFC| 1.62×
SD of LFC 1.57×
fraction |LFC| > 1 4.00×
max |LFC| 1.41×
Negative control 1.00×
Cliff's δ (memory vs matched control, 95% CI)
c
Donor 1
Donor 2
Donor 3
Donor 4
% Higher median response
+35%
+47%
+66%
+51%
CORAL
GEMLI
Power-Seek
ClonePair

**Supplementary Fig. 3: Memory-gene response is robust to the expression floor, the response metric, and the donor.**
**a** Availability of expression-matched control genes, per screen. Genes were sorted by mean control expression and cut into 50 equal-count bins. Grey, the number of background genes available in each bin, that is, genes no method placed in its top 300; red, the number of memory genes in that bin requiring a match. The count axis is logarithmic. Annotations give the tightest region of each screen, above the 95th percentile of memory-gene expression, as the number of candidates remaining for the memory genes there.
**b** Effect size under different expression floors and different metrics to measure response, for genes called by at least two of the four methods in the T cell screen (Zhu). Points, Cliff's delta between memory genes and expression-matched controls; bars, bootstrap 95% confidence intervals (300 resamples); right column, the ratio of median response in memory genes to that in controls. Controls are matched to the memory genes on mean control expression (50 quantile bins, averaged over 100 draws). Top, the floor on mean control expression, applied before the gene universe is defined, from no floor to 2 UMI per control cell; bold, the floor used in all other analyses (0.1 UMI per control cell; leaving 335 memory genes in a universe of 5,734). Bottom, the statistic summarising each gene's response across perturbations, all at the 0.1 UMI floor: mean |LFC| (bold, used in all other analyses), median |LFC|, standard deviation of the LFC, fraction of perturbations with |LFC| > 1, and maximum |LFC|. Grey, a negative control, the number of perturbations in which each gene was measured, which reflects sampling depth rather than response and is therefore expected not to differ between memory genes and controls.
**c** Response computed separately within each of the four donors of the T cell screen (Zhu), per method (top 300 genes per method). Points, each donor's percentage increase in median response of memory genes over expression-matched controls, one colour per donor; bars, the 95% confidence interval of the mean across donors (t distribution, n = 4 donors); numbers, that mean; dashed line, no difference. The interval reflects variability between donors rather than the number of genes tested.

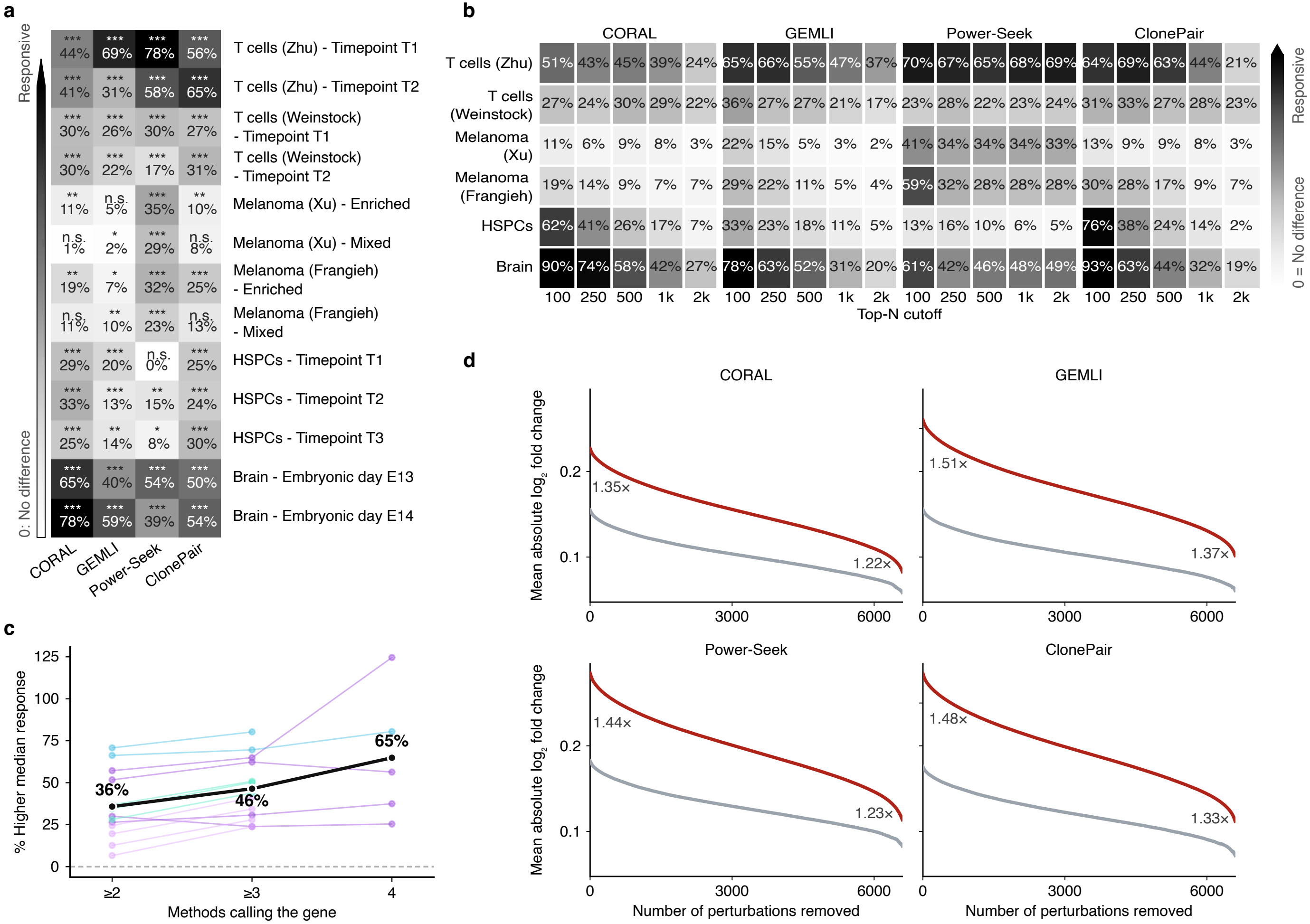

a
Responsive
0: No difference
*** 44% *** 69% *** 78% *** 56% T cells (Zhu) - Timepoint T1
*** 41% *** 31% *** 58% *** 65% T cells (Zhu) - Timepoint T2
*** 30% *** 26% *** 30% *** 27% T cells (Weinstock) - Timepoint T1
*** 30% *** 22% *** 17% *** 31% T cells (Weinstock) - Timepoint T2
** 11% n.s. 5% *** 35% ** 10% Melanoma (Xu) - Enriched
n.s. 1% * 2% *** 29% n.s. 8% Melanoma (Xu) - Mixed
** 19% * 7% *** 32% *** 25% Melanoma (Frangieh) - Enriched
n.s. 11% ** 10% *** 23% n.s. 13% Melanoma (Frangieh) - Mixed
*** 29% *** 20% n.s. 0% *** 25% HSPCs - Timepoint T1
*** 33% *** 13% ** 15% *** 24% HSPCs - Timepoint T2
*** 25% ** 14% * 8% *** 30% HSPCs - Timepoint T3
*** 65% *** 40% *** 54% *** 50% Brain - Embryonic day E13
*** 78% *** 59% *** 39% *** 54% Brain - Embryonic day E14
CORAL GEMLI Power-Seek ClonePair
b
CORAL GEMLI Power-Seek ClonePair
T cells (Zhu) 51% 43% 45% 39% 24% 65% 66% 55% 47% 37% 70% 67% 65% 68% 69% 64% 69% 63% 44% 21%
T cells (Weinstock) 27% 24% 30% 29% 22% 36% 27% 27% 21% 17% 23% 28% 22% 23% 24% 31% 33% 27% 28% 23%
Melanoma (Xu) 11% 6% 9% 8% 3% 22% 15% 5% 3% 2% 41% 34% 34% 34% 33% 13% 9% 9% 8% 3%
Melanoma (Frangieh) 19% 14% 9% 7% 7% 29% 22% 11% 5% 4% 59% 32% 28% 28% 28% 30% 28% 17% 9% 7%
HSPCs 62% 41% 26% 17% 7% 33% 23% 18% 11% 5% 13% 16% 10% 6% 5% 76% 38% 24% 14% 2%
Brain 90% 74% 58% 42% 27% 78% 63% 52% 31% 20% 61% 42% 46% 48% 49% 93% 63% 44% 32% 19%
100 250 500 1k 2k
Top-N cutoff
Responsive
0 = No difference
c
% Higher median response
125 100 75 50 25 0
36% 46% 65%
≥2 ≥3 4
Methods calling the gene
d
CORAL GEMLI Power-Seek ClonePair
Mean absolute log2 fold change
1.35× 1.22× 1.51× 1.37× 1.44× 1.23× 1.48× 1.33×
0.2 0.1
0 3000 6000
Number of perturbations removed

**Supplementary Fig. 4: The response effect holds in every experimental group, grows with method agreement, does not depend on the top-N cutoff, and is not driven by only a few perturbations.**

**a** Increase in median response of memory genes over matched controls, as in **Fig. 2g**, with every lineage dataset group entering as its own row. Rows, screen and group; columns, method; memory genes are each method's top 300 within that group; colour, the percentage increase, with zero marking no difference.

**b** Increase in median response of memory genes over expression-matched controls, for every lineage dataset and screen pairing (rows), every method (column blocks) and every top-N cutoff (columns within a block: 100, 250, 500, 1,000, 2,000). Memory genes are each method's top N; controls are the genes of no method placed in the same top N, matched on mean control expression (50 equal-count bins, averaged over 100 draws). Colour, the percentage increase, with zero marking no difference. Where a method ranks fewer than N genes it contributes all of them. For those methods the memory set stops growing with N while the background, defined as the genes no method placed in its top N, continues to shrink, so the deeper columns compare the same gene set against a progressively more restricted control pool.

**c** The same quantity as in **a,** with memory genes defined by increasing method agreement, as in **Fig. 2h**: genes ranked in the top 300 by at least two, at least three or all four methods within a group. One line per screen-by-group; black, the mean across combinations.

**d** Response after removing the most impactful perturbations, for each method separately. The 6,646 perturbations of the T cell screen (Zhu[27]) were ranked by their mean absolute $\log_2$ fold change across that method's top 300 memory genes and removed one at a time, strongest first. Red, mean absolute log fold change (LFC) of the memory genes over the perturbations that remain; grey, the same for an expression-matched set of genes that no method called. Ratios between the two curves are given at the start and end of each panel.

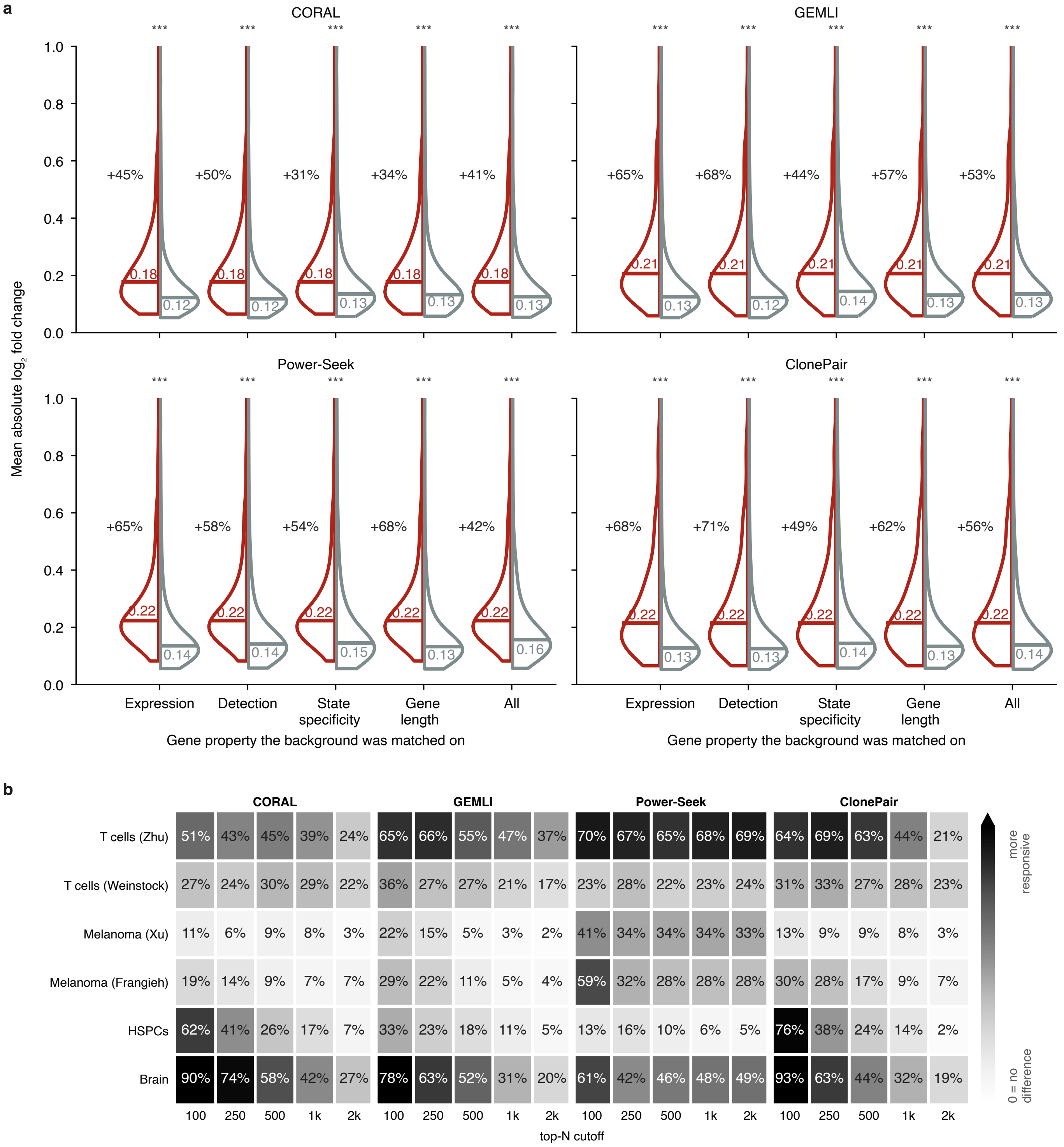
a
CORAL
GEMLI
Power-Seek
ClonePair
Mean absolute $\log_2$ fold change
***
+45%
+50%
+31%
+34%
+41%
+65%
+68%
+44%
+57%
+53%
+65%
+58%
+54%
+68%
+42%
+68%
+71%
+49%
+62%
+56%
Expression
Detection
State specificity
Gene length
All
Gene property the background was matched on
b
CORAL
GEMLI
Power-Seek
ClonePair
T cells (Zhu) 51% 43% 45% 39% 24% 65% 66% 55% 47% 37% 70% 67% 65% 68% 69% 64% 69% 63% 44% 21%
T cells (Weinstock) 27% 24% 30% 29% 22% 36% 27% 27% 21% 17% 23% 28% 22% 23% 24% 31% 33% 27% 28% 23%
Melanoma (Xu) 11% 6% 9% 8% 3% 22% 15% 5% 3% 2% 41% 34% 34% 34% 33% 13% 9% 9% 8% 3%
Melanoma (Frangieh) 19% 14% 9% 7% 7% 29% 22% 11% 5% 4% 59% 32% 28% 28% 28% 30% 28% 17% 9% 7%
HSPCs 62% 41% 26% 17% 7% 33% 23% 18% 11% 5% 13% 16% 10% 6% 5% 76% 38% 24% 14% 2%
Brain 90% 74% 58% 42% 27% 78% 63% 52% 31% 20% 61% 42% 46% 48% 49% 93% 63% 44% 32% 19%
100 250 500 1k 2k
top-N cutoff
more responsive
0 = no difference

**Supplementary Fig. 5: Memory genes experimentally identified from bulk RNA-seq by MemorySeq significantly overlap those determined computationally from lineage-barcoded datasets and show the same elevated response.**

**a** Overlap between lineage-barcoded memory genes and bulk MemorySeq genes in melanoma. Harmange[18] memory genes, taken as each method's top 300 within each experimental group (population enriched for primed state, mixed population of all live cells; **Methods**), are compared with the 227 MemorySeq hits of Shaffer et al.[16] in WM989 cells. Bars, the number of shared genes at each level of method agreement; solid red, enriched; hatched red, mixed; grey, an expression-matched baseline in which the memory set is redrawn within its own expression-rank bins (20 bins, 1,000 permutations), with bar height the permutation mean and whisker the 97.5th percentile.

**b** Response of both gene sets in the two melanoma screens. Mean absolute $\log_2$ fold change per gene across all perturbations of the Xu[31] and Frangieh[32] screens, for Harmange lineage-barcoded memory genes (called by at least two of the four methods within their top 300, enriched and mixed pooled) and for the Shaffer bulk MemorySeq genes, each against its own expression-matched background (50 bins, averaged over 100 draws). Red, memory genes; grey, matched background; numbers inside each half, medians; percentages, the increase in median over the background; asterisks, two-sided Mann-Whitney U test with Benjamini-Hochberg correction across the two comparisons of each screen (** $q < 0.01$, *** $q < 0.001$).

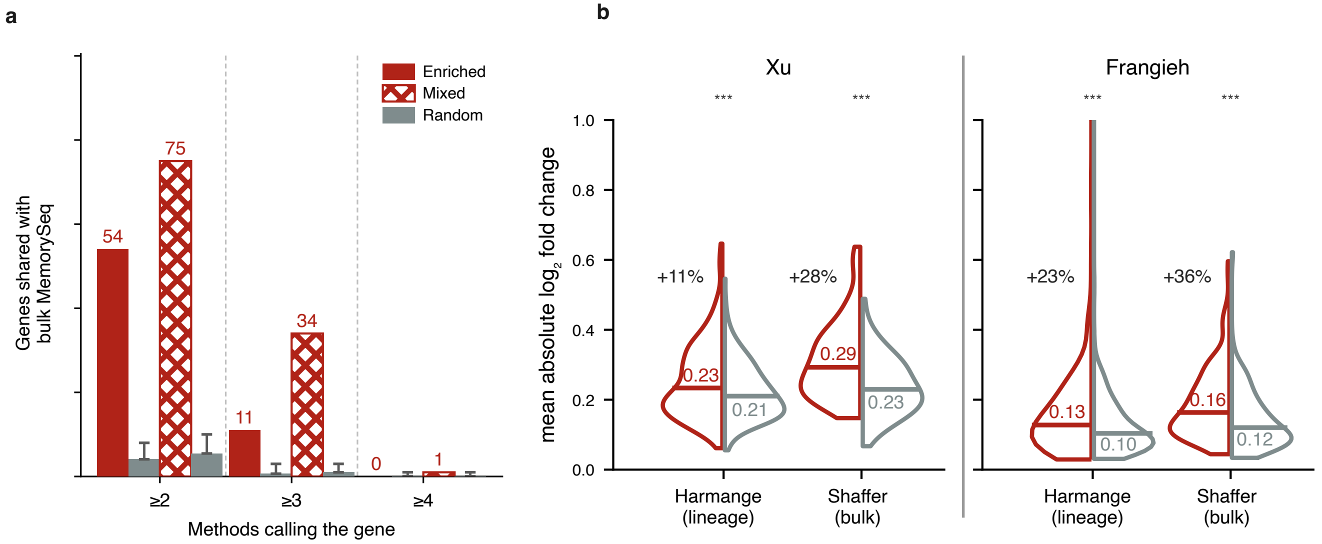
a
Enriched
Mixed
Random
Genes shared with bulk MemorySeq
54
75
11
34
0
1
≥2
≥3
≥4
Methods calling the gene
b
Xu
Frangieh
***
***
***
***
mean absolute $\log_2$ fold change
1.0
0.8
0.6
0.4
0.2
0.0
+11%
+28%
+23%
+36%
0.23
0.21
0.29
0.23
0.13
0.10
0.16
0.12
Harmange (lineage)
Shaffer (bulk)
Harmange (lineage)
Shaffer (bulk)

**Supplementary Fig. 6: Memory genes directly identified in control cells of perturbation screens without lineage barcodes show the same elevated response.**

**a** Response of memory genes defined without lineage information, one row per Perturb-seq screen. Within a row, each gene set is compared with its own expression-matched background: red, memory genes; grey, background. Three callers are shown, each in two versions. Control cells (bold labels): the caller was run on that screen's own unperturbed control cells with no lineage information. Power-Seek[21] requires none by design, GEMLI[20] uses clones predicted from expression, and isCORAL[40] uses pseudoclones inferred from expression. Barcode: the same caller was run on the cell-type-matched lineage dataset, with CORAL[15] as isCORAL's barcoded counterpart. For GEMLI and CORAL, this means using the experimental clone labels; Power-Seek does not use clone labels in either column, so for it the two columns compare the screen's control cells with the lineage dataset's cells. The last column is the set called by all three methods on the control cells. All sets are the top 300 genes of their ranking, compared with background matched on mean control expression (50 equal-count bins, averaged over 100 draws) among genes above 0.1 UMI per control cell. Horizontal lines inside each half, medians; percentages, the increase in median over the matched background; asterisks, two-sided Mann-Whitney U test with Benjamini-Hochberg correction across the seven gene sets of each screen (n.s., $q > 0.05$; * $q < 0.05$; ** $q < 0.01$; *** $q < 0.001$). Where a set contains fewer genes than the others, its size is given beneath the label.

**b** The same estimates as in **a** on one axis. Each marker is one gene set from **a**, positioned by the percentage by which its median response exceeds its own expression-matched background; the dashed line, no difference. Rows, screens; colour, caller; filled circles, the control-cell set; open circles, the barcode-derived set, so a segment pointing right means the barcode-free route gave the more responsive set. Dark diamonds, the set called by all three methods on the control cells. Statistics as in **a**.

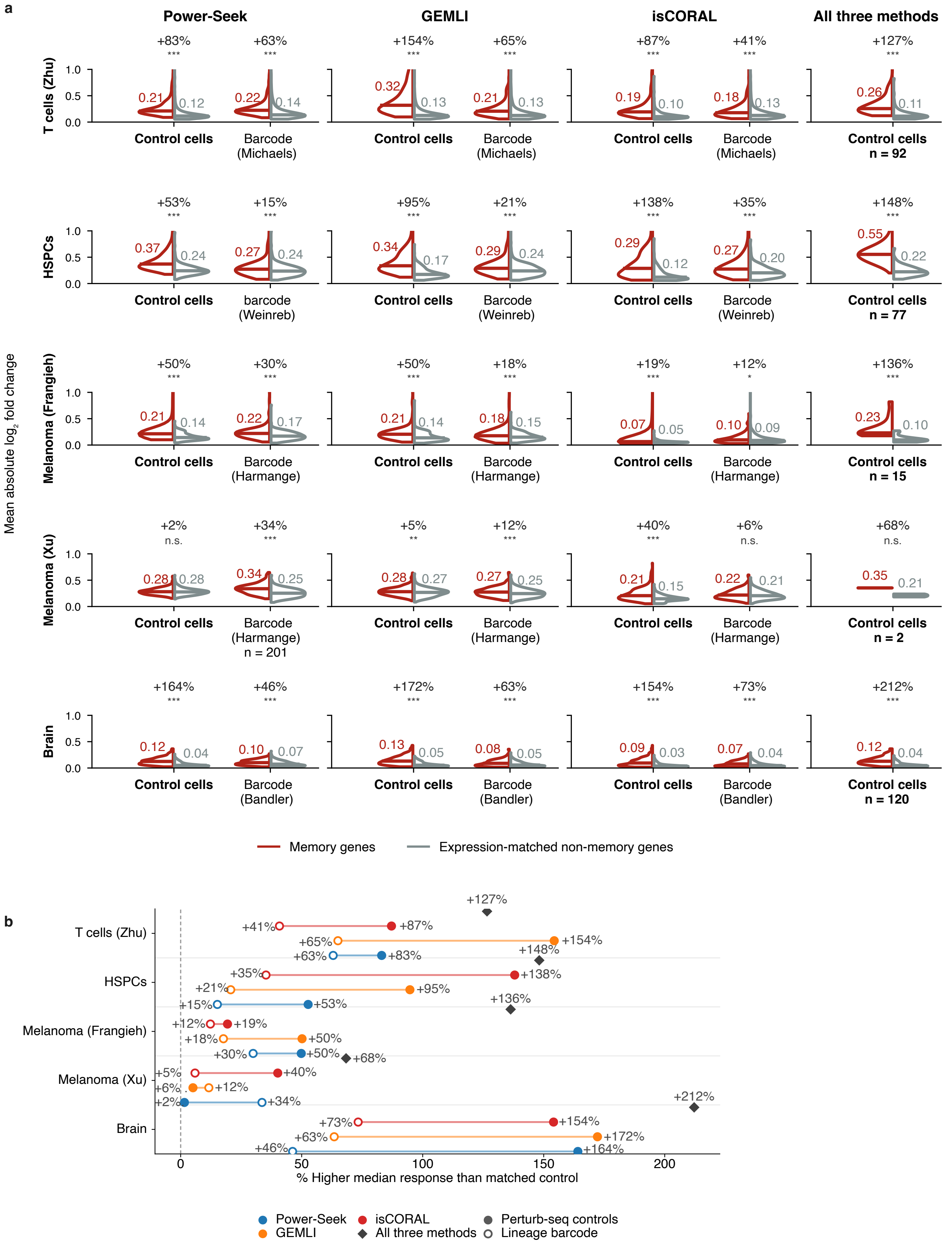
a
Power-Seek
GEMLI
isCORAL
All three methods
T cells (Zhu)
HSPCs
Melanoma (Frangieh)
Melanoma (Xu)
Brain
Mean absolute log2 fold change
1.0
0.5
0.0
+83% *** +63% *** +154% *** +65% *** +87% *** +41% *** +127% ***
0.21 0.12 0.22 0.14 0.32 0.13 0.21 0.13 0.19 0.10 0.18 0.13 0.26 0.11
Control cells
Barcode (Michaels)
Control cells n = 92
+53% *** +15% *** +95% *** +21% *** +138% *** +35% *** +148% ***
0.37 0.24 0.27 0.24 0.34 0.17 0.29 0.24 0.29 0.12 0.27 0.20 0.55 0.22
barcode (Weinreb)
Barcode (Weinreb)
Control cells n = 77
+50% *** +30% *** +50% *** +18% *** +19% *** +12% * +136% ***
0.21 0.14 0.22 0.17 0.21 0.14 0.18 0.15 0.07 0.05 0.10 0.09 0.23 0.10
Barcode (Harmange)
Control cells n = 15
+2% n.s. +34% *** +5% ** +12% *** +40% *** +6% n.s. +68% n.s.
0.28 0.28 0.34 0.25 0.28 0.27 0.27 0.25 0.21 0.15 0.22 0.21 0.35 0.21
Barcode (Harmange) n = 201
Control cells n = 2
+164% *** +46% *** +172% *** +63% *** +154% *** +73% *** +212% ***
0.12 0.04 0.10 0.07 0.13 0.05 0.08 0.05 0.09 0.03 0.07 0.04 0.12 0.04
Barcode (Bandler)
Control cells n = 120
Memory genes
Expression-matched non-memory genes
b
T cells (Zhu)
HSPCs
Melanoma (Frangieh)
Melanoma (Xu)
Brain
+127% +41% +87% +65% +154% +63% +83%
+148% +35% +138% +21% +95% +15% +53%
+136% +12% +19% +18% +50% +30% +50% +68%
+5% +40% +6% +12% +2% +34%
+212% +73% +154% +63% +172% +46% +164%
0
50
100
150
200
% Higher median response than matched control
Power-Seek
GEMLI
isCORAL
All three methods
Perturb-seq controls
Lineage barcode

**Supplementary Fig. 7: Memory genes identified without lineage information are more responsive in every Perturb-seq screen tested.**

Power-Seek[21] was run on each screen's own unperturbed control cells, without lineage barcodes or clone labels, and its top 300 genes were compared with an equal-sized set of genes it did not call, matched on control expression (50 equal-count bins, averaged over 100 draws). Red, memory genes; grey, matched background; point, medians across genes; connector, the two medians of one screen. The response is the mean absolute $\log_2$ fold change of a gene across all perturbations of that screen, with each perturbation's own target masked, computed over genes above 0.1 UMI in the controls. Above each pair, the difference between the two medians and a two-sided Mann-Whitney U test, Benjamini-Hochberg corrected across all 18 comparisons (**$q < 0.01$, ***$q < 0.001$). Screens are ordered by that difference. To the right of the dashed line, the three screens that also contain paired perturbations are shown a second time, scored on those pairs alone; their single-perturbation comparisons use the same memory genes and appear in the main block. Screen labels give the study and the cell line or condition profiled.

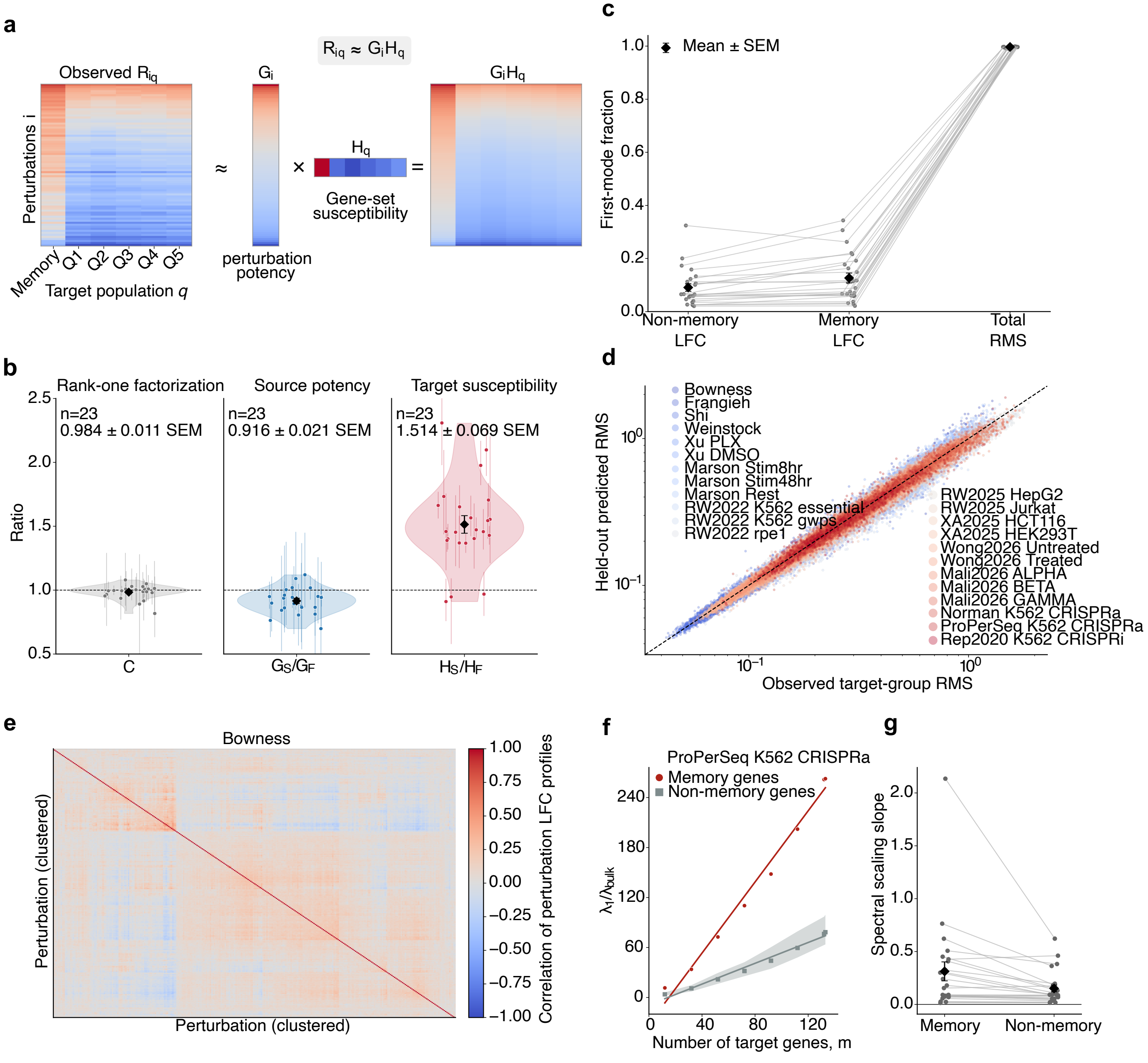
a
Observed $R_{iq}$
$R_{iq} \approx G_i H_q$
$G_i$
$G_i H_q$
Perturbations i
≈
×
$H_q$
=
Gene-set susceptibility
perturbation potency
Memory
Q1
Q2
Q3
Q4
Q5
Target population q
b
Rank-one factorization
Source potency
Target susceptibility
n=23
0.984 ± 0.011 SEM
n=23
0.916 ± 0.021 SEM
n=23
1.514 ± 0.069 SEM
Ratio
C
$G_S/G_F$
$H_S/H_F$
c
Mean ± SEM
First-mode fraction
Non-memory LFC
Memory LFC
Total RMS
d
Bowness
Frangieh
Shi
Weinstock
Xu PLX
Xu DMSO
Marson Stim8hr
Marson Stim48hr
Marson Rest
RW2022 K562 essential
RW2022 K562 gwps
RW2022 rpe1
RW2025 HepG2
RW2025 Jurkat
XA2025 HCT116
XA2025 HEK293T
Wong2026 Untreated
Wong2026 Treated
Mali2026 ALPHA
Mali2026 BETA
Mali2026 GAMMA
Norman K562 CRISPRa
ProPerSeq K562 CRISPRa
Rep2020 K562 CRISPRi
Held-out predicted RMS
Observed target-group RMS
e
Bowness
Perturbation (clustered)
Correlation of perturbation LFC profiles
f
ProPerSeq K562 CRISPRa
Memory genes
Non-memory genes
$\lambda_1/\lambda_{bulk}$
Number of target genes, m
g
Spectral scaling slope
Memory
Non-memory

**Supplementary Fig. 8: Memory genes are more susceptible targets, but not more potent sources of perturbation.**

**a** Schematic of the separable response model. The observed matrix of root-mean-square (RMS) responses $R_{iq}$, with rows indexing perturbations i and columns indexing target gene populations q (memory genes and non-memory quintiles Q1-Q5 ordered by mRNA half-life), is approximated by the product of a perturbation potency $G_i$ and a gene-set susceptibility $H_q$.

**b** Factorization constant C (gray), source-potency ratio $G_S/G_F$ (blue) and target-susceptibility ratio $H_S/H_F$ (red), where S and F denote memory and non-memory classes. Perturbations and measured genes in each dataset were split into these classes and the pooled RMS of signed log fold changes (LFCs) computed for each of the four source-target blocks; C, $G_S/G_F$ and $H_S/H_F$ were derived from the resulting 2x2 amplitude matrix (**SI**). Unequal group sizes were controlled by 300 bootstrap draws per dataset, resampled to the smaller source- and target-group sizes; dataset-level estimates are bootstrap medians and light vertical lines are 2.5th-97.5th bootstrap percentiles. Translucent violins show the distribution across datasets, colored points individual datasets, and black diamonds the across-dataset mean ± SEM (n = 23 datasets with sufficiently represented classes). Dashed lines mark unity; dataset-level values were tested against unity by two-sided one-sample t-tests.

**c** Fraction of squared response amplitude captured by the leading mode $f_1$ of a singular-value decomposition. For gene-level matrices (perturbations x target genes, signed log-fold changes, columns centered across perturbations), $f_1$ is shown for memory genes and for matched non-memory controls (median over 20 control sets matched without replacement by proximity in log RNA half-life). For total RMS, $f_1$ was computed from an uncentered, strictly positive perturbation-by-population matrix with six columns (memory genes and non-memory quintiles Q1-Q5), so that a large value reflects common-amplitude, not necessarily common-direction structure. Thin gray lines join the three values within a dataset; gray points are individual datasets (horizontally jittered) and black diamonds the mean ± SEM (n = 24).

**d** Four-fold cross-validation of the rank-one prediction. Within each dataset, geometric-mean ratios between the RMS response of each gene group and that of reference group Q1 were fitted on three folds of perturbations and used to predict the response of each held-out perturbation from its measured Q1 response. Each point is one held-out perturbation and one gene group (memory genes and Q2-Q5; Q1 excluded as the reference), colors denote datasets, and the dashed line is the identity (454,150 predictions across 24 datasets; median $R^2 = 0.964$ in log space).

**e** Pearson correlation between the signed LFC response profiles of pairs of perturbations in the Bowness dataset, hierarchically clustered by average linkage on the distance 1 − correlation. Off-diagonal block structure indicates that distinct perturbations elicit recurring, correlated response programs.

**f** Ratio of the leading to the median bulk eigenvalue, $\lambda_1/\lambda_{bulk}$, of the gene-gene response correlation matrix as a function of the number of target genes m, in ProPerSeq K562 CRISPRa. Memory genes (red) and matched non-memory controls (grey; median of 49 independently sampled control sets matched one-to-one by RNA half-life) are shown with fitted lines $\lambda_1(m)/\lambda_{bulk}(m) = b + sm$. Independent responses give a ratio of approximately one at all m, whereas a coherent shared component gives $1 + \alpha m$, so the slope s measures the strength of shared response structure.

**g** Spectral scaling slopes s for memory genes and matched non-memory controls across 24 datasets. Gray points are individual datasets, connecting lines preserve the pairing, and black diamonds show the across-dataset mean ± SEM (two-sided paired t-test, $n = 24$, $p = 0.021$).
